# The phenuivirus Toscana virus makes an atypical use of vacuolar acidity to enter host cells

**DOI:** 10.1101/2023.03.06.531240

**Authors:** Jana Koch, Qilin Xin, Martin Obr, Alicia Schäfer, Nina Rolfs, Holda Anagho, Aiste Kudulyte, Lea Woltereck, Susann Kummer, Joaquin Campos, Zina M. Uckeley, Lesley Bell-Sakyi, Hans-Georg Kräusslich, Florian KM Schur, Claudio Acuna, Pierre-Yves Lozach

## Abstract

Toscana virus is a major cause of arboviral disease in humans in the Mediterranean basin during summer. However, early virus-host cell interactions and entry mechanisms remain poorly characterized. Investigating iPSC-derived human neurons and cell lines, we found that virus binding to the cell surface was specific but inefficient, and 50% of bound virions were endocytosed within 10 min. Virions entered Rab5a+ early endosomes and, subsequently, Rab7a+ and LAMP-1+ late endosomal compartments. Penetration required intact late endosomes and occurred within 30 min following internalization. Virus entry relied on vacuolar acidification, with an optimal pH for viral membrane fusion at pH 5.5. The pH threshold increased to 5.8 with longer pre-exposure of virions to the slightly acidic pH in early endosomes. Strikingly, the particles remained infectious after entering late endosomes with a pH below the fusion threshold. Overall, our study establishes Toscana virus as a late-penetrating virus and reveals an atypical use of vacuolar acidity by this virus to enter host cells.

## Introduction

Toscana virus (TOSV) is a re-emerging sand fly-borne pathogen of the *Phenuiviridae* family (genus *Phlebovirus*, order *Bunyavirales*) that is responsible for neuro-invasive infections in humans and causes meningitis and meningoencephalitis in most severe cases (*1, 2*). The virus was first isolated from the phlebotomine sand flies *Phlebotomus perniciosus* and *Phlebotomus perfiliewi* in Tuscany, central Italy, in 1971 (*3*). Nowadays, TOSV is widely spread in North Africa and southern Europe, including Greece, Italy, southern France and Spain (*1, 2, 4*). TOSV is currently a primary cause of arthropod-borne viral disease in humans in Mediterranean countries during summer (*2*). However, until now, no vaccines or antiviral treatments are approved for human use.

TOSV has a tri-segmented, single-stranded RNA genome of predominantly negative polarity that replicates exclusively in the cytosol of infected cells (*5*). The larger segment (L) codes for the RNA-dependent RNA polymerase that is required to initiate virus replication after release of the viral genome into the cytosol. The medium segment (M) encodes a polyprotein precursor, the proteolytic cleavage of which results in a non-structural protein, NSm, and two envelope glycoproteins, Gn and Gc. The smallest genomic segment (S) codes for the non-structural protein NSs, and for the nucleoprotein N which associates with the RNA genome and constitutes, together with the viral polymerase, the ribonucleoproteins (RNPs) (*6*). Viral particles are believed to assemble and acquire their lipid envelope in the endoplasmic reticulum or Golgi network, from where newly-formed viral particles leave infected cells.

TOSV has so far not been visualized, and hence the morphology, size and structural organization of virions remain to be determined. Other phenuiviruses are enveloped, roughly spherical, and about 100 nm in diameter (*7*). While viral RNPs are inside the viral particles, the two envelope glycoproteins Gn and Gc decorate the outer surface and allow virus binding to host cells and acid-activated penetration into the cytosol. Cryo-electron tomography revealed that most regular phenuiviral particles have protrusions of about ten nanometers forming an icosahedral lattice with an atypical T=12 triangulation (*8, 9*). Structural studies of Rift Valley fever virus (RVFV), Dabie virus (DABV) and Heartland virus (HRTV) revealed that the phenuiviral Gc belongs to the group of class-II membrane fusion proteins (*10–12*).

The tropism, receptors, cellular factors and pathways used by TOSV to enter and infect host cells are largely unidentified and poorly characterized. The virus was shown to subvert heparan sulfates and the C-type lectins DC-SIGN and L-SIGN to attach to the cell surface (*13–15*). A few phenuiviruses have been shown to depend on endocytic internalization and vacuolar acidification for infectious entry (*7*). As with other class-II fusion proteins, acidic pH is thought to trigger multiple conformational changes in phenuiviral glycoproteins that lead to the insertion of the viral fusion unit into endosomal membranes (*16*). Foldback of the viral fusion protein follows and then the formation of a fusion pore allows the release of the virus genome into the cytosol.

Here, we analyzed the entry of TOSV into induced pluripotent stem cell (iPSC)-derived human neurons and other tissue culture cells. To this end, we developed sensitive fluorescence-based approaches to examine and quantify TOSV infection, binding, internalization, intracellular trafficking and membrane fusion. The results showed that TOSV shares with other phenuiviruses a dependence on the degradative branch of the endocytic machinery for penetration of host cells by acid-activated membrane fusion. TOSV made atypical use of endosomal acidity to find its way through endosomal vesicles and enter the cytosol.

## Results

### TOSV infects human iPSC-derived neurons

TOSV causes meningoencephalitis in the most severe cases. Therefore, we sought to test the sensitivity of brain cells to TOSV infection. To this end, functional, human glutamatergic neurons were generated from iPSCs through expression of the transcription factor neurogenin-2 (*17*) and exposed to different multiplicities of infection (MOIs) of TOSV for 48 h. The susceptibility of cells was assessed by flow cytometric analysis after immunofluorescence staining with antibodies (Abs) directed against all TOSV structural proteins, *i.e.*, the nucleoprotein N and the glycoproteins Gn and Gc. Nearly 70% of the neuronal cells were infected at the highest MOI (Fig. 1A). The percentage of infected cells increased over time and reached a plateau within 16 h post-infection (hpi) (Fig. 1B), indicating that the fluorescence signal detected in this assay corresponded to viral replication and not to the input virions.

**Fig. 1.**
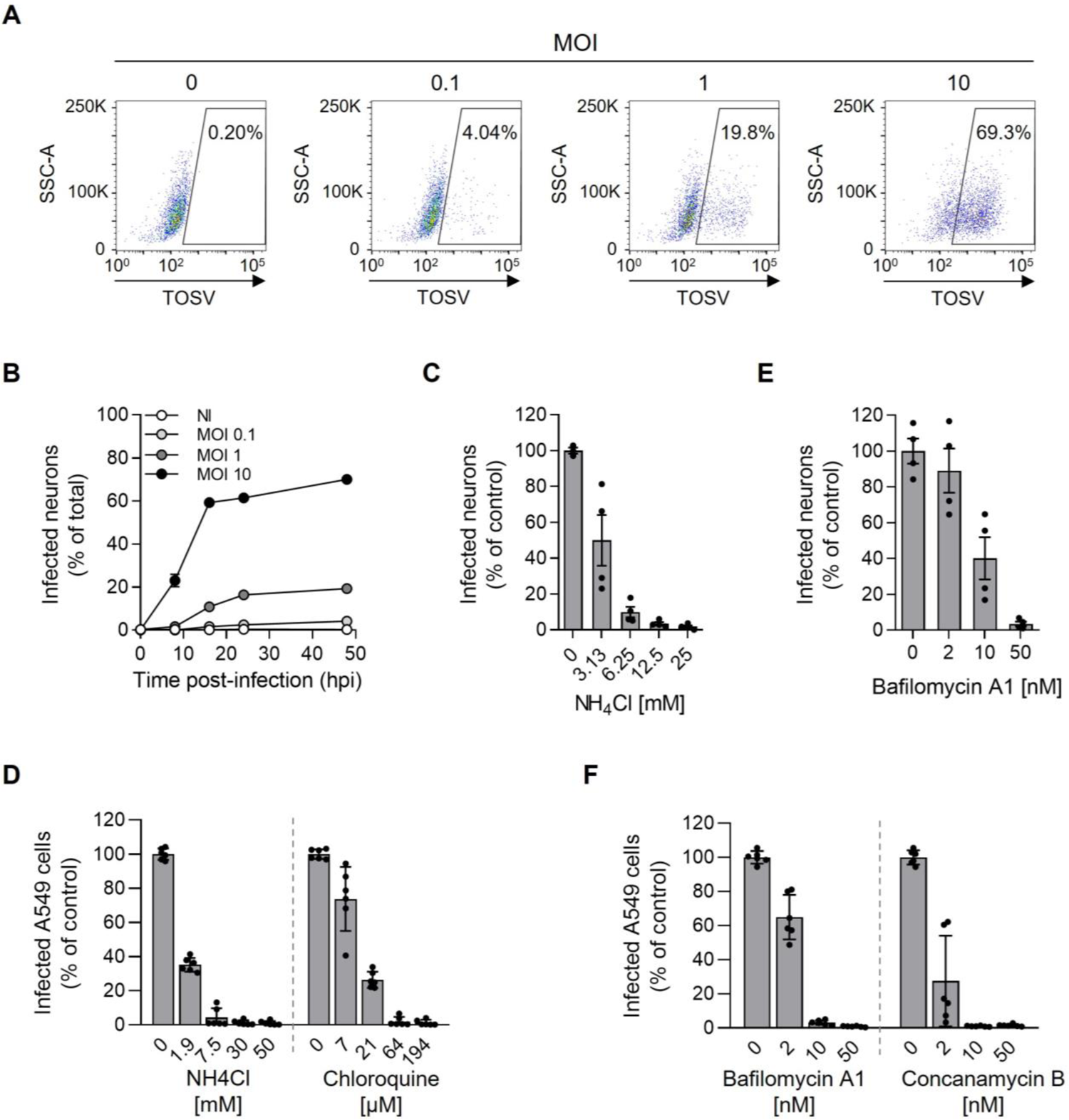
Toscana virus (TOSV) entry into iPSC-derived human neurons depends on endosomal acidification. (**A**) Induced human pluripotent stem cell (iPSC)-derived neurons were exposed to TOSV at the indicated multiplicities of infection (MOIs) and harvested 48 h later. After fixation and permeabilization, infected cells were stained with a polyclonal antibody against the TOSV structural proteins N, Gn and Gc. Infection was then quantified by flow cytometry. (**B**) iPSC-derived neurons were infected with TOSV at the indicated MOIs, and infection was monitored over 48 h using the flow cytometry-based assay described in panel A. (**C** to **F**) iPSC neurons (C and E) and A549 cells (D and F) were pretreated with agents that elevate endosomal pH at the indicated concentrations and were infected with TOSV at MOI 10 for 8 h and MOI 2 for 6 h, respectively, in the continuous presence of ammonium chloride (NH_4_Cl) (C and D), chloroquine (D), bafilomycin A1 (E and F) and concanamycin B (F). Infection was analyzed by flow cytometry as described in Fig. 1A. Data were normalized to those of control samples without inhibitor treatment.

Several cell types of various species have been reported to support productive TOSV infection (*5, 14*), suggesting broad host range potential and wide tissue tropism of the virus. We evaluated 18 further cell lines from different arthropod and vertebrate species and found that 16 were susceptible and permissive to infection and seven of eight tested cell lines produced infectious viral particles, as determined by our flow cytometric and plaque-forming unit (pfu) assays (Table S1 and Fig. S1A and S1B). Of note, myeloid or lymphatic lineages were poorly infected, if at all. The three sand fly cell lines allowed a complete viral cycle, but their sensitivity to virus infection was low compared to that of most of the mammalian cells.

To evaluate the production and release of infectious viral particles, we infected A549 cells at low MOIs and quantified virus infection and production up to 48 hpi (Fig. S1C to S1E). Infectious progeny was found to be released from infected cells as early as 9-16 hpi. Collectively, our analysis indicated that TOSV completes one round of infection, from virus binding and penetration to replication and release of infectious progeny within 9 h in A549 cells. Similar results were obtained in other cell lines tested (Table S1). As we aimed to analyze TOSV entry mechanisms and restrict infection to a single round in the selected cell lines, we limited our assays to 6 hpi in all further experiments. In addition, we used MOIs for each cell line allowing the infection of approximately 20% of cells. This range of infection generally avoids saturation of infection in cells and thus, allows the detection of potential inhibitory or enhancing effects of a perturbant.

### TOSV infection relies on vacuolar acidification

To examine whether the acidic pH in endosomal vesicles is important in TOSV infection, the virus was added to neurons and A549 cells in the presence of agents that neutralize vacuolar pH. The lysosomotropic weak bases ammonium chloride (NH_4_Cl) and chloroquine induced a dose-dependent inhibition of TOSV infection (Fig. 1C and 1D). In these experiments, we monitored and regulated the medium for a pH above 7.0 ensuring effectiveness of the base. Two inhibitors of vacuolar-type H+-ATPases, bafilomycin A1 and concanamycin B, gave similar results (Fig. 1E and 1F). Taken together, these experiments showed that TOSV depends on vacuolar acidification for infection in both neurons and A549 cells. These results also suggest that TOSV enters host cells by endocytosis, as reported for the other phenuiviruses analyzed for entry (*7*).

Together, this series of data indicates that TOSV efficiently infects human neurons and various mammalian cell lines. Regardless of cell type, infection required endosomal acidification. To further examine TOSV entry and the role of endosomal maturation and vacuolar acidification, we, therefore, selected three productive cell lines representing rodent, monkey and human cells, namely BHK-21, Vero and A549 cells. In most of the following experiments, we opted to use these cell lines as they are easier to handle than the complex iPSC-derived neuron system.

### Labeling of TOSV with fluorescent dyes

To visualize and quantify the early stages of TOSV entry, we purified and labeled the virus chemically with fluorescent probes. Hydroxysuccinimidyl (NHS) ester dyes with an excitation wavelength of 488 nm (Alexa Fluor [AF] 488) or 647 nm (ATTO647N) were coupled to free lysines in the glycoproteins Gn and Gc at a dye-to-glycoprotein ratio between 1:1 and 2:1. At this ratio, we assumed that all virions were labeled. Alternatively, TOSV was labeled with the lipid dye R18 primarily for the analysis of viral fusion. A high concentration of R18 molecules in viral envelopes results in autoquenching of the fluorescence signal (*18*). Viral fusion allows the release of R18 into the target cell membranes leading to the dilution of the dye, and, thus, dequenching of the fluorescence.

Characterization of the fluorescent virions is shown in Figure S2. Briefly, labeled particles were purified through a sucrose gradient so that unbound dyes were removed (Fig. S2A). Analysis by SDS-PAGE and Coomassie blue staining showed that the purity of the labeled TOSV preparations was greater than 90% (Fig. S2B). The only proteins labeled with NHS ester dyes were Gn and Gc (Fig. S2B). The observation that N was not labeled demonstrated that the viral envelope was intact during the labeling procedure. The different labeled particles could be visualized as single spots by confocal microscopy and super-resolution stimulated emission depletion (STED) microscopy (Fig. S2C). We did not notice significant impact of labeling on TOSV infectivity. The titers were similar to those of non-labeled particles (Fig. S2D).

TOSV particles were then imaged by cryo-electron microscopy (EM) after fixation with paraformaldehyde and vitrification (Fig. 2A). Virions appeared roughly spherical with a diameter of 121 ± 11 nm (n = 96) and protrusions of 9 ± 2 nm (n = 96) (Fig. 2B). The measured roundness coefficient of virions was close to 1, *i.e.*, the ratio between their perpendicular width and length was of 0.9 ± 0.1 (n = 96) (Fig. 2B). This reflected the nearly spherical shape of the viral particles. Overall, TOSV particles displayed the typical morphology known for other phenuiviruses such as RVFV and Uukuniemi virus (UUKV) (*8, 9*).

**Fig. 2.**
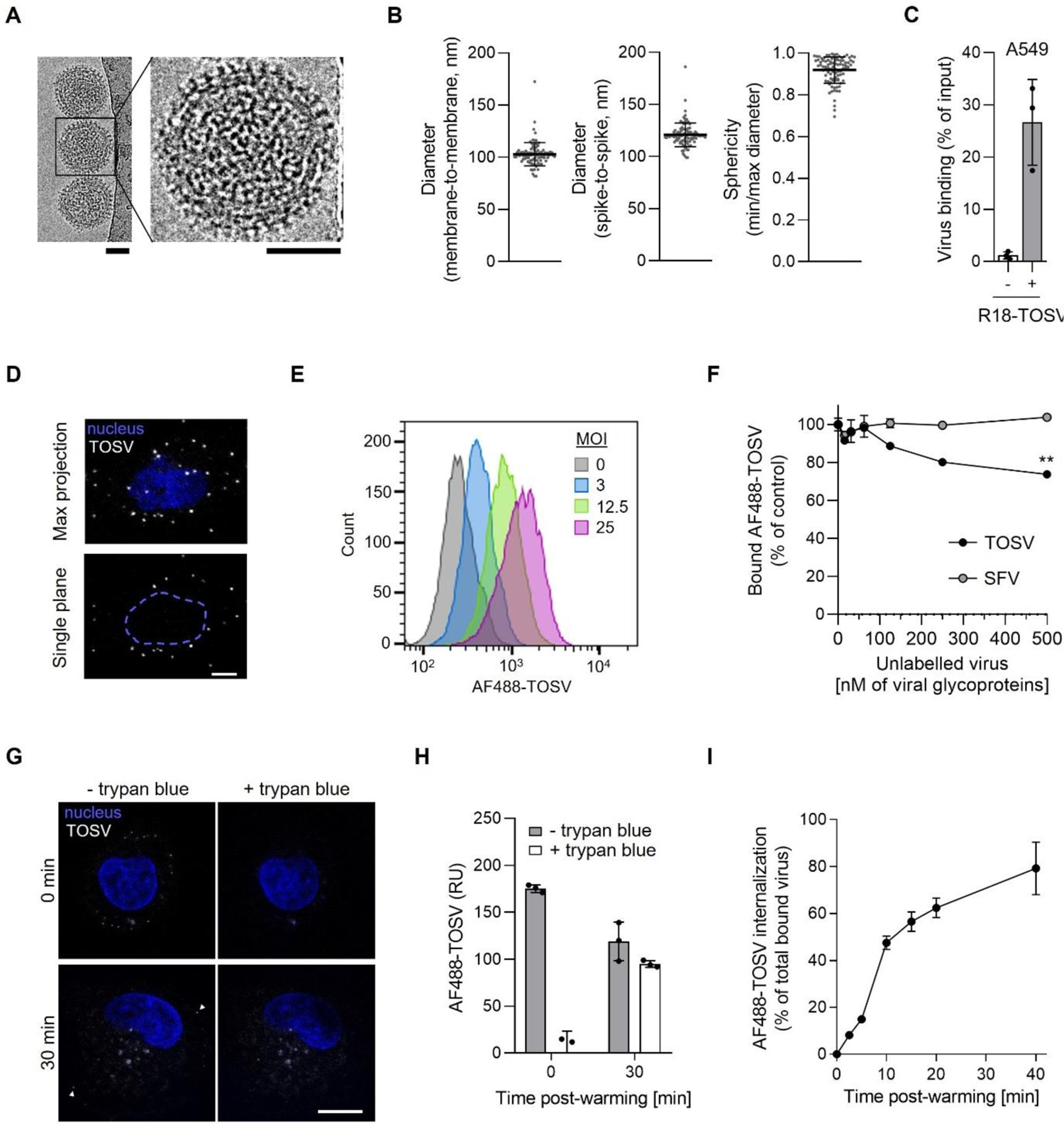
Binding of Toscana virus (TOSV) to human A549 cells is inefficient and internalization slow. (**A**) TOSV virions were purified, fixed with paraformaldehyde, vitrified and imaged by cryo-electron microscopy (EM). Scale bar, 50 nm. (**B**) The membrane-to-membrane and spike-to-spike diameter of virions was measured from EM images. The sphericity of each TOSV particle was determined as the ratio of the width to the perpendicular length (n = 96). (**C**) R18-labeled TOSV (+) or no virus (-) was bound to A549 cells on ice at MOI 10 for 90 min. Virus binding was analyzed with a spectrofluorometer after extensive washing of the cells to remove unbound viruses. The virus input was measured in cell-free suspension before binding. The data correspond to the fraction of the virus input bound to cells, *i.e.*, the ratio of fluorescence associated with cells following virus binding to that measured in the virus input. (**D**) ATTO647N-labeled TOSV was bound to A549 cells at MOI 1 on ice, fixed, and nuclei were stained with Hoechst before imaging with a confocal microscope. The upper panel shows cell-associated virions (white spots) seen in maximum z-projection images acquired in the 647-nm channel, while the bottom image shows one focal plane with virions in white and nuclei indicated by a blue dashed line. Scale bar, 5 µm. (**E**) AF488-labeled TOSV (AF488-TOSV) was bound to A549 cells on ice at the indicated MOIs for 1.5 h before fixation and analysis by flow cytometry. (**F**) A549 cells in suspension were first exposed to increasing amounts of unlabeled TOSV and Semliki Forest virus (SFV) on ice for 45 min and then to AF488-TOSV (15 nM of glycoproteins) for an additional hour on ice. Cells were washed and virus binding was measured by flow cytometry. Data were normalized to those in cells not pre-exposed to the unlabeled virus. T-test with Welch’s correction was applied. **, p<0.01. (**G**) AF488-TOSV (white) was bound to A549 cells on ice at MOI 10 for 1.5 h before warming to 37°C for 30 min. Cells were then washed, fixed and treated with trypan blue before confocal imaging. Nuclei were stained with Hoechst (blue). Arrowheads show some fluorescent particles at the cell surface that are quenched upon trypan blue addition. Scale bar, 10 µm. (**H**) AF488-TOSV was bound to A549 cells in suspension at MOI 10 at 4 °C before warming to 37 °C for 30 min. Cells were treated with trypan blue before analysis by flow cytometry. RU, relative unit. (**I**) AF488-TOSV was bound to A549 cells in suspension at 4 °C and rapidly warmed to 37 °C to allow virus uptake for up to 40 min. Endocytic internalization of virions was determined by flow cytometry after trypan blue treatment. Internalization is given as the percentage of fluorescence quantified in samples treated with trypan blue compared to that in untreated samples. The fluorescence signal measured in cells not exposed to AF488-TOSV was considered as the background signal and subtracted from the other values.

### TOSV binding to cells is specific but rather inefficient

To evaluate TOSV binding to cells, R18-labeled virions were first allowed to bind to A549 cells in suspension at 4 °C for 90 min. The unbound virions were washed away, and the remaining cell-associated, fluorescent viral particles were determined with a fluorimeter. The results showed that the fraction associated with the cells was about 25% of the virus input (Fig. 2C), indicating rather inefficient binding of TOSV to cells. After binding of ATTO647N-labeled particles to A549 cells at 4 °C, virus particles were imaged by confocal microscopy and could be detected on the cell surface (Fig. 2D). That the spots had varied sizes suggested that not only individual virions were attached to cells. The largest clusters were probably formed by 2-3 virions but were only seen as a single spot due to the limitation of confocal resolution. The fact that the number of spots per cell was 10.5 ± 8.5 (n = 151) at MOI only 1, *i.e.*, one infectious virion initially added per cell, indicated that the ratio between infectious and noninfectious bound virions was around 1:10.

Flow cytometry analysis allowed detection and quantification of TOSV-AF488 from MOI 3 and above (Fig. 2E). Binding of TOSV-AF488 was observed to be abrogated by pre-binding of non-labeled TOSV in a dose-dependent manner (Fig. 2F). A 33-fold higher concentration of non-labeled virions reduced TOSV-AF488 binding by one-fourth. Complete inhibition was not achieved because the necessary concentrations of non-labeled virus were not reached under our experimental settings. However, the pre-binding of Semliki Forest virus (SFV) did not affect TOSV-AF488 binding under the same conditions. Combined, these data indicated that TOSV binding to cells is relatively inefficient and likely involves one or more specific attachment factors or receptors.

### TOSV enters early and late endosomal compartments

We next aimed to determine whether TOSV is internalized and sorted into early endosomes (EEs) following virus binding to the cell surface. To analyze TOSV internalization by fluorescence microscopy, we first allowed TOSV-AF488 to bind to A549 cells on ice at high MOI (∼10). The cells were then rapidly warmed to 37 °C to trigger endocytosis and placed back on ice after 30 min to stop further endocytosis. To discriminate between internalized and surface-bound virions, cells were treated with trypan blue for 10 sec before imaging or flow cytometry analysis. Trypan blue is a membrane-impermeable dye that quenches green-emitting dyes such as AF488 and thus quenches the fluorescence emitted by TOSV-AF488 particles exposed on the cell surface while leaving intracellular virions unquenched (Fig. 2G and 2H). Time-course analysis of the generation of trypan blue-resistant fluorescence of cell-associated TOSV-AF488 revealed that internalization into A549 cells started within the first 5 min and increased over time to reach the half-maximal level (t_1/2_) within 9 ± 2 min and the plateau 10 min later (Fig. 2I). Evidently, TOSV uptake occurred rather synchronously.

To assess whether internalization leads to the sorting of viral particles into EEs, we used A549 cells transiently expressing the small GTPase Rab5a, a marker of EEs, tagged with a monomeric enhanced green fluorescent protein (EGFP). After synchronization of TOSV-ATTO647N binding to A549 cells expressing EGFP-Rab5a on ice, the temperature was rapidly shifted to 37 °C for periods of up to 40 min. Confocal microscopy showed TOSV co-localizing with EGFP-Rab5a-positive (+) vesicles 5 min post-warming (Fig. 3A). The amount of co-localizing virions reached a maximum within 5-10 min post-warming and decreased thereafter (Fig. 3B). In live A549 cells, confocal microscopy showed that TOSV-ATTO647N moved together within EGFP-Rab5a+ endosomal vacuoles (Movie S1).

**Fig. 3:**
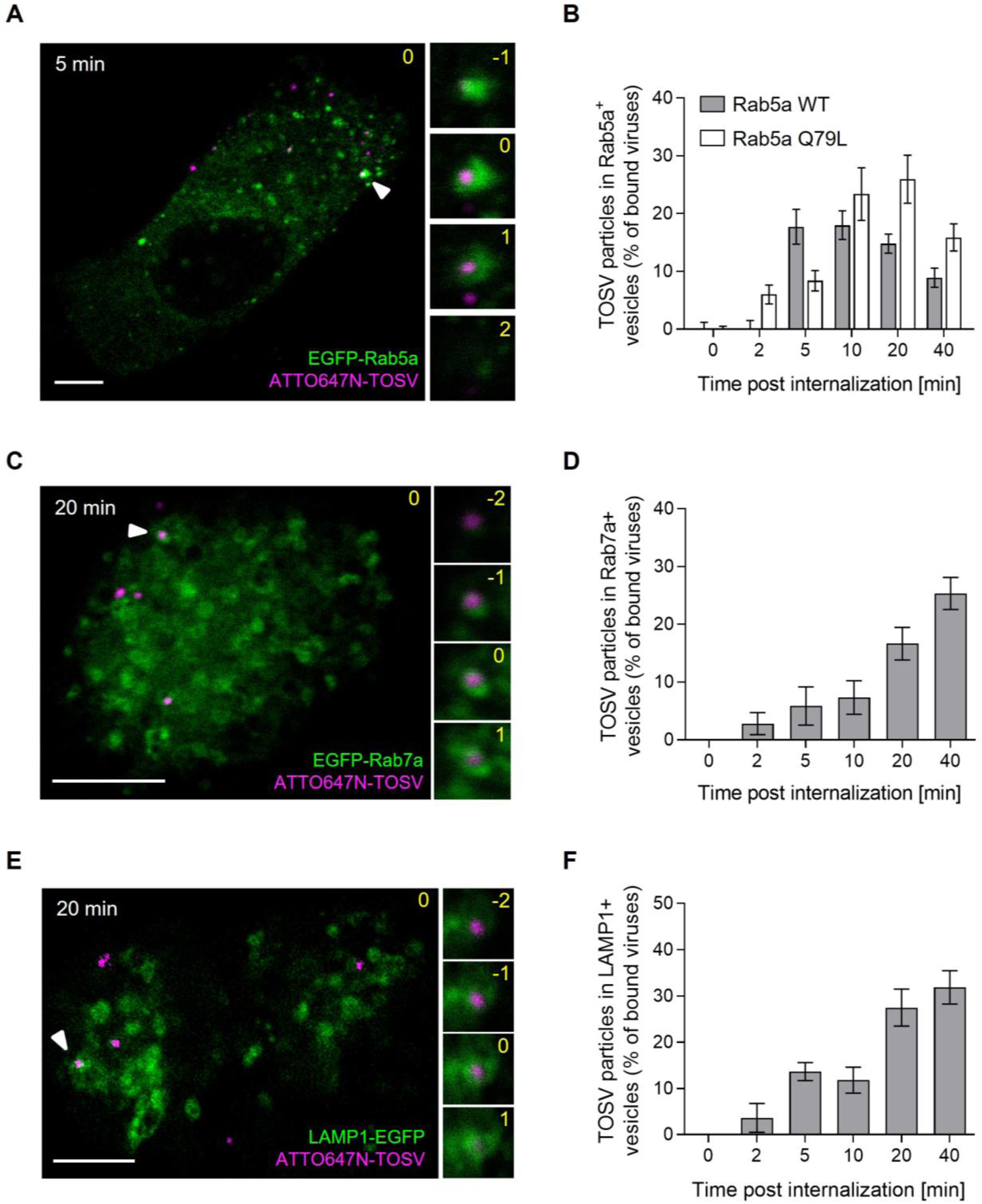
Toscana virus (TOSV) enters early endosomes and then late endosomal organelles. (**A**) A549 cells transiently expressing EGFP-Rab5a were exposed to ATTO647N-TOSV at MOI 1 for 90 min on ice and then rapidly warmed for 5 min at 37 °C to allow the internalization of virions. Cells were subsequently washed and fixed. TOSV (magenta) and Rab5a (green) were imaged by confocal microscopy. One focal plane is shown. Higher magnifications of association between TOSV and Rab5a-positive (+) vesicles (white arrowhead) are shown on the right side as a z-stack series. Yellow numbers indicate the position of the stack in the series, and the original plane is marked with 0. Scale bar, 5 µm. (**B**) Internalization of ATTO647N-TOSV was monitored in cells transiently expressing EGFP-Rab5a wild-type (WT) or its constitutively active mutant Q79L over a time span of 40 min as described in A. Co-localization is expressed as the percentage of bound TOSV associated with Rab5a+ vesicles at different times post-warming. A minimum of 6 cells were analyzed per time point. (**C** to **F**) Prebound ATTO647N-TOSV was internalized into A549 cells transiently expressing EGFP-Rab7a (C and D) or LAMP1-EGFP (E and F) for up to 40 min before confocal microscopy and image-based quantification as described in A and B. At least 9 cells were analyzed per condition. Scale bar, 5 µm.

In addition, TOSV was observed within vesicles containing the late endosomal markers Rab7a and LAMP-1 tagged with EGFP (EGFP-Rab7a and LAMP-1-EGFP) but at later time points. Co-localization and coordinated motion with EGFP-Rab7a+ endosomes in live A549 cells were maximal 20-40 min after uptake (Fig. 3C and 3D and Movie S2). Co-localization with the lysosome marker LAMP-1 (LAMP-1-EGFP) was somewhat delayed (Fig. 3E and 3F). Of note, some virions were located in the middle of the vesicles and had probably not yet undergone fusion with the limiting endosomal membrane. Overall, the temporal overlap of co-localizations with Rab7 and LAMP-1 suggested that viral particles reach late endosomes (LEs) rather than lysosomes.

### TOSV depends on the passage through EEs and LEs for infectivity

To examine whether passage through the endosomal compartments was required for infectivity, we first assessed TOSV internalization and infection in A549 cells transfected with DNA plasmids to express a constitutively-active mutant of Rab5a tagged with EGFP (EGFP-Rab5a Q79L). The expression of this mutant typically results in the enlargement of EEs (Fig. 4A), compromising the maturation of LEs and transport of cargo to lysosomes (*19*). Infection was measured in populations of cells expressing identical levels of EGFP as selected by flow cytometry. Expression of EGFP-Rab5a Q79L reduced TOSV infection by 80% in comparison to EGFP-Rab5a wild type (wt) (Fig. 4B). In contrast to EGFP-Rab5a, the number of virions co-localizing with EGFP-Rab5a Q79L+ vacuoles remained high even after 20-40 min (Fig. 3B). Expression of the dominant-negative mutant Rab5a S34N (EGFP-Rab5a S34N), which abrogates the maturation of newly-formed EEs (*20*), also resulted in a large decrease in infection, *i.e.*, more than 60% (Fig. 4B). Altogether, these results indicated that the infectious entry pathway involves the passage of TOSV in Rab5a+ EEs, though the transport to downstream endosomal vesicles is also needed for productive infection.

**Fig. 4:**
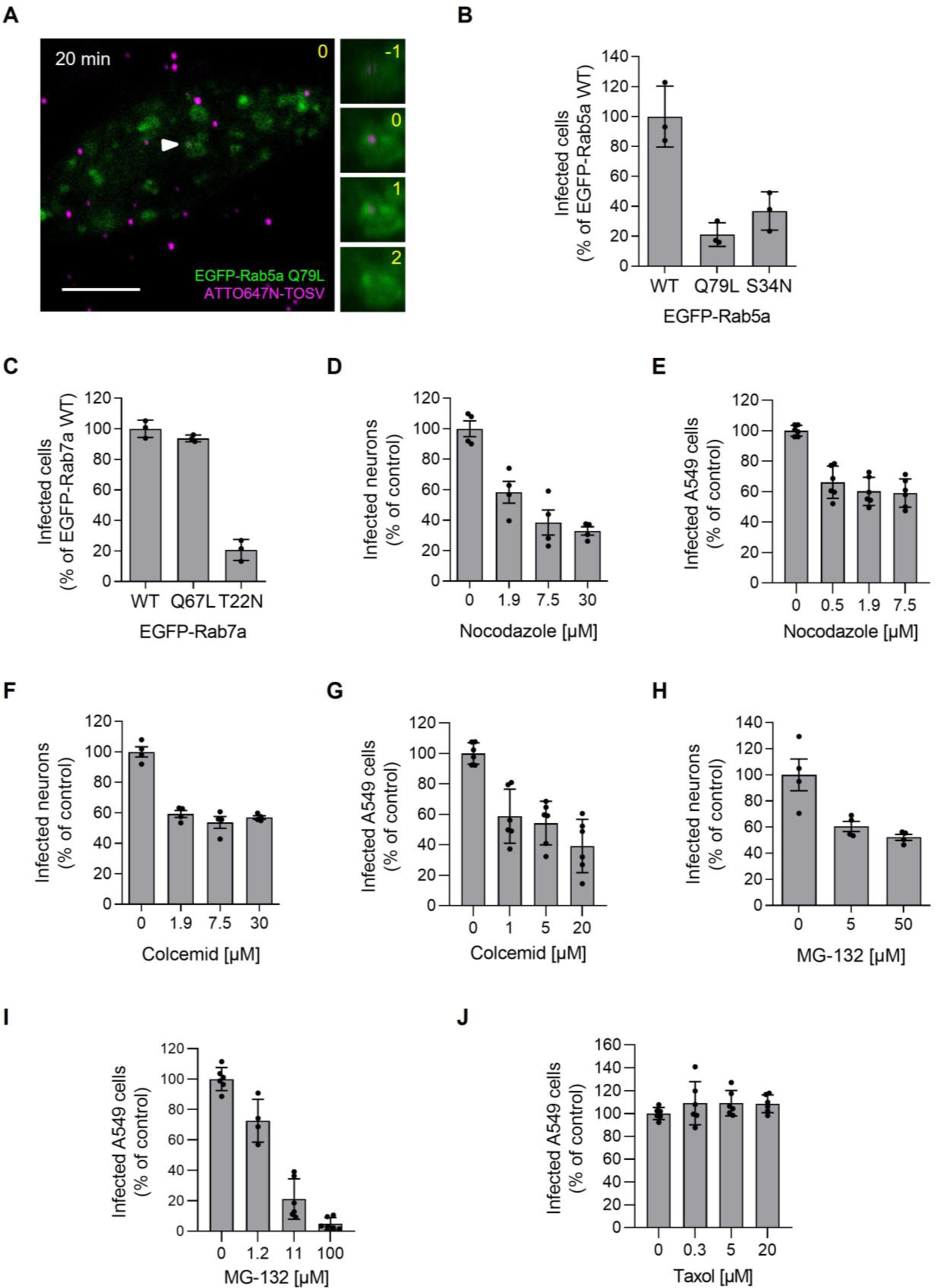
Toscana virus (TOSV) relies on late endosomal maturation for infectious entry. (A) ATTO647N-TOSV at MOI 1 was allowed to enter A549 cells transiently expressing the constitutively active mutant Q79L of EGFP-Rab5a as described in Fig. 3. Internalization of virions was allowed for 20 min at 37 °C, and cells imaged by confocal microscopy. TOSV is seen in magenta and EEs containing EGFP-Rab5a Q79L are in green. Scale bar, 5 µm. (**B**) EGFP-Rab5 WT, its constitutively active mutant Q79L and its dominant-negative mutant S34N were transiently expressed in A549 cells. Transfected cells were then challenged with TOSV at MOI 4 for 6 h. Cell populations with a similar level of Rab5a expression were selected with a flow cytometer, and each population was analyzed for infection. Infection was normalized to that in the cell population expressing EGFP-Rab5 WT. (**C**) A549 cells transiently expressing EGFP-Rab7a WT, its constitutively active mutant Q67L, or its dominant-negative mutant T22N were challenged with TOSV for 6 h before flow cytometry analysis following the approach used in the panel B. (**D** to **J**) iPSC-derived neurons (D, F and H) and A549 cells (E, G, I and J) were pretreated with nocodazole (D and E), colcemid (F and G), MG-132 (H and I), or taxol (J) at indicated concentrations, and then infected with TOSV at MOI 10 (iPSC neurons) and 2 (A549) in the continuous presence of inhibitors for 8 h and 6 h, respectively. Infection was quantified by flow cytometry, and data were normalized to those in control samples without inhibitor treatment.

As a block in the maturation of EEs to LEs impeded infection, TOSV potentially needs to pass through LEs for its productive entry into the cytosol. To test this possibility, we evaluate the role of the small GTPase Rab7a in TOSV infection. Rab7a is a key player in LE maturation and functions (*21*). When A549 cells transiently expressed an EGFP-tagged dominant negative mutant of Rab7a (EGFP-Rab7a T22N), TOSV infection was severely impaired, but not when the cells expressed the constitutively active Q67L mutant of Rab7a (Fig. 4C). In some cell types, LE maturation relies on microtubule (MT)-mediated transport of endosomes to the perinuclear region of cells and on free cytosolic ubiquitin (*22, 23*). Treatment of neurons and A549 cells with either nocodazole or colcemid, two drugs that hamper MT polymerization, resulted in a 40-60% decrease in infection (Fig. 4D to 4G). When free ubiquitin was depleted by the proteasome inhibitor MG-132, TOSV infection was reduced in a dose-dependent manner in both neurons and A549 cells (Fig. 4H and 4I). Conversely, taxol, an MT-stabilizing drug, did not affect the infection of A549 cells (Fig. 4J). Taken together, these experiments show that the transport of virions to LEs is required for infectivity.

### Low pH is sufficient and necessary for TOSV fusion

The observation that TOSV requires vacuolar acidification for infection suggests that the virus penetrates host cells by acid-activated membrane fusion. To define the pH threshold and to link fusion with infection, we tested the capacity of TOSV to fuse at the plasma membrane of cells as originally described for SFV (*24*). In such a scenario, the virus bypasses the need for endocytosis during productive infection. Briefly, TOSV was bound to A549 cells at MOI 10 at 4 °C, the temperature was rapidly elevated to 37 °C for 1.5 min in buffers with different pH values, and NH_4_Cl-containing medium at neutral pH was then added for the remaining period of infection to prevent further infection through endosomes. The bypass resulted in efficient infection at pH values of 5.7 and below (Fig. 5A). 50% of the maximal infection was reached at a pH of 5.6. These data demonstrated that a reduction in pH is sufficient to trigger infectious penetration of viral RNPs from the plasma membrane to the cytosol. Additional processing of the viral glycoproteins Gn and Gc in the endocytic machinery were apparently not needed to activate fusion and infection by TOSV.

**Fig. 5:**
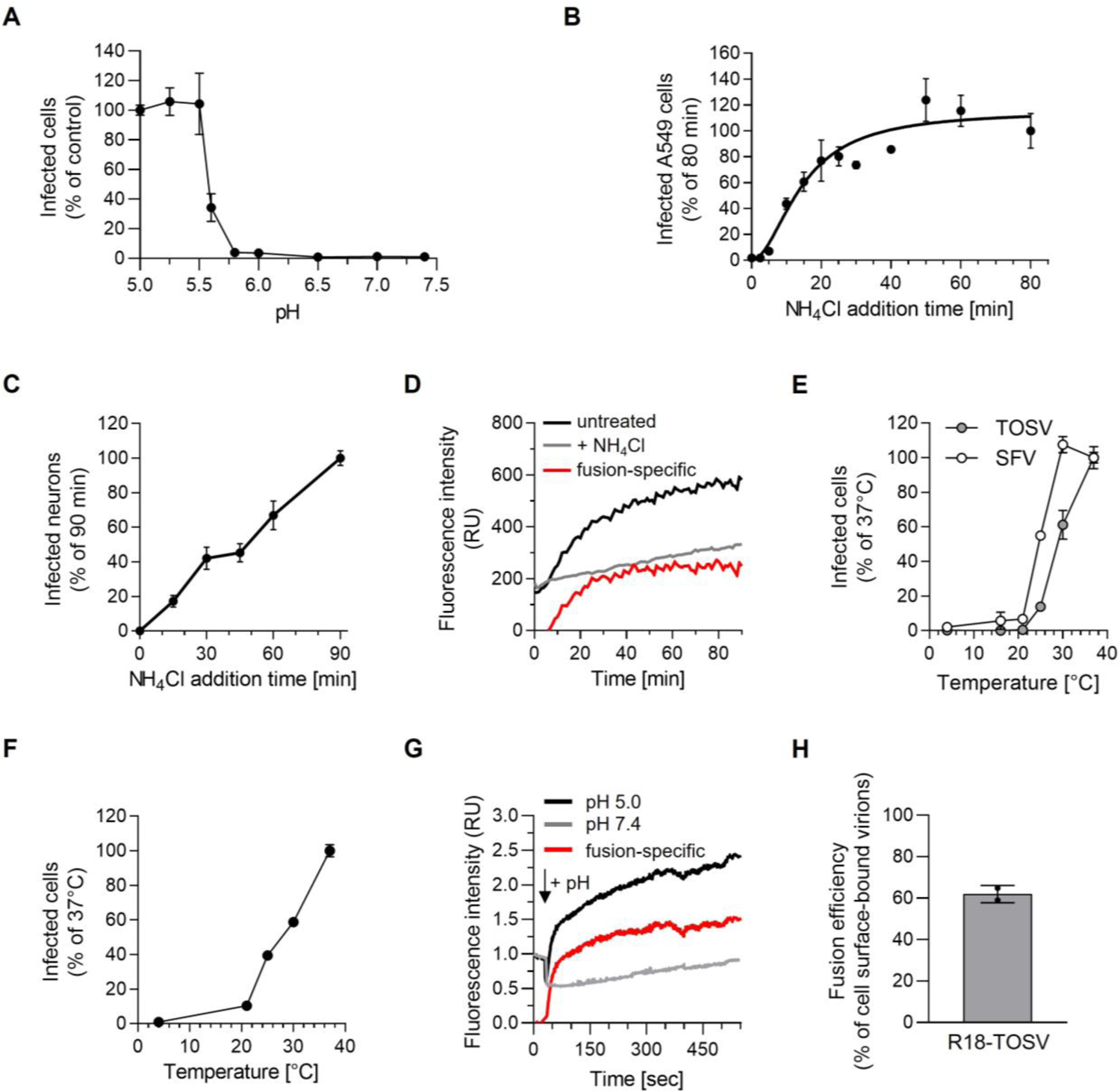
Toscana virus (TOSV) penetrates host cells by acid-activated membrane fusion. (**A**) TOSV was bound at MOI 10 to A549 cells for 1.5 h on ice. Subsequently, cells were washed and subjected to the indicated pH values at 37 °C for 90 sec to trigger the virus fusion at the plasma membrane. Infected cells were then incubated for 7 h at 37 °C in the presence of 50 mM NH_4_Cl to prevent viral penetration from endosomes, and thereby only monitor the release of viral genomes from the plasma membrane. Infection was quantified by flow cytometry, and the data were normalized to those from samples where the infection was triggered with a buffer at pH 5.0. (**B** and **C**) TOSV particles were bound to A549 cells (B) and iPSC-derived neurons (C) at MOIs 1 and 15, respectively, on ice for 90 min and then rapidly shifted to 37 °C to allow virus internalization. 50 mM NH_4_Cl was added at the indicated times to block further viral penetration. Infected cells were quantified by flow cytometry, and data were normalized to the samples where NH_4_Cl was added 80 min (B) and 90 min (C) post-warming, respectively. (**D**) R18-TOSV was bound at MOI 10 to A549 cells on ice and rapidly warmed to 37 °C for 90 min. The increase in fluorescence resulted from the dequenching of the lipid dye R18 after virus fusion with cell membranes in living cells and was measured with a spectrofluorometer. NH_4_Cl was used to block virus fusion by neutralizing endosomal pH and, thus, to define the fluorescence background due to spontaneous translocation of the R18 dyes between the viral envelope and the neighboring cell membrane (grey line). The red line shows the virus fusion-specific R18 release, *i.e.*, black line (fusion + free diffusion) minus grey line (free diffusion). RU, relative unit. (**E**) TOSV and Semliki Forest virus (SFV) were bound to A549 cells on ice, and samples were shifted to indicated temperatures for 50 min. Infected cells were subsequently incubated at 37°C for 6 h in the presence of NH_4_Cl to block further viral penetration. Infection was analyzed by flow cytometry and normalized to that in samples incubated throughout at 37 °C. (**F**) TOSV fusion efficiency was assessed at the indicated temperatures using the plasma membrane-virus fusion assay described in panel A. Data were normalized to those of samples incubated throughout at 37 °C. (**G**) R18-TOSV binding to A549 cells at MOI 10 was synchronized on ice for 90 min. Infected cells were then placed in a spectrofluorometer, the temperature warmed to 37 °C, and the fluorescence signal was monitored over 550 sec. “+pH” indicates when a buffer at pH 5.0 (black line) or 7.4 (grey line) was added to the samples to trigger virus fusion at the plasma membrane, *i.e.*, 30 sec later, once the temperature reached 37 °C. Data were normalized to those at the time point 0. The red line shows the virus fusion-specific R18 release at pH ∼5.0, *i.e.*, black line (fusion at pH ∼5.0 + free diffusion) minus grey line (free diffusion at pH ∼7.4). RU, relative unit. (**H**) The protocol described in D was used to record real-time penetration of R18-TOSV into A549 cells at MOI 10 for 90 min. Triton X-100 was then added to the samples to induce the dequenching of all R18 molecules, allowing the measure of total fluorescence in the cells that correlated with all bound and internalized virions. The data show the fraction of plasma membrane-bound viruses that had fused in the cells, *i.e.*, the ratio of the fluorescence associated with the viral fusion to that associated with all virions in the cells.

### Acid-activated penetration occurs in late endosomal compartments

To determine the timing of the acid-requiring step following virus internalization, we took advantage of the fact that the rise in endosomal pH is almost instantaneous when NH_4_Cl is added to the extracellular medium (*25*). Virus particles were first allowed to bind to neurons and A549 cells on ice at MOIs of 15 and 1, respectively. Virus entry was then synchronized by rapid warming to 37 °C, and NH_4_Cl was added at different times following the temperature switch. A concentration of 50 mM was used to ensure that infection was completely abolished after adding NH_4_Cl. Infectious penetration started after a 5-min delay in A549 cells and reached a t_1/2_ within 15 min and a plateau 25-45 min later (Fig. 5B). It was apparent that individual viral particles had completed the NH_4_Cl-sensitive step non-synchronously in neurons, most likely due to the heterogeneity of cell preparations typical of iPSC-derived neurons (Fig. 5C).

To further analyze acid-activated membrane fusion in late endosomal vacuoles, we relied on TOSV-R18 to monitor viral fusion in living cells with a fluorimeter. In this assay, the increase in fluorescence signal results from dequenching of the fluorescence lipid dye R18 upon activation of viral fusion. Though the subsequent release of the dye into the target cell membranes corresponds to the hemi-fusion mixture of outer leaflets and not to fusion pore formation, it is a good correlate for fusion. TOSV-R18 was bound to A549 cells on ice, and virus endocytosis was synchronized by switching the cells rapidly to 37 °C (Fig. 5D). The kinetics measured in this assay were very similar to those determined with the above procedure based on NH_4_Cl addition. The fluorescence signal started to increase after a 6 min lag and reached a t_1/2_ within 18 min ± 2 min post-warming with a plateau value about 20 min later.

The kinetics of penetration closely resembled the time course of endolysosomal maturation, which usually lasts 30-60 min (*22*). To challenge the hypothesis that TOSV penetration occurs in LEs, we examined the temperature dependence of entry. The transport of cargo from EEs to LEs is known to be inhibited at temperatures below 20 °C (*26*). TOSV binding to A549 cells was synchronized on ice, and cells rapidly warmed to different temperatures for 1 h before incubation at 37 °C for 6 h in the presence of NH_4_Cl to prevent further penetration. The infection was greatly reduced at 30 °C and below (Fig. 5E). In contrast to TOSV, a temperature of 30 °C had no noticeable effect on infection by SFV, which is acid-activated in EEs (*27*). SFV infection was still detected at temperatures as low as 16 °C.

To rule out that the fusion process was altered by temperature, we also analyzed the temperature dependence of TOSV fusion using the bypass protocol described above. At 25°C, fusion corresponded to 40% of the 37 °C control, whereas infection via the normal route was lowered by 85% (Fig. 5E and 5F). At 21 °C, fusion was still 10% of the 37 °C control, whereas infection via the normal route could no longer be detected. As expected of a virus capable of replicating in insect hosts, this suggested that fusion was not a bottleneck for penetration at lower temperatures. Most likely the viral particles did not infect at 25 °C and below because they did not reach a compartment with low enough pH.

Next, we analyzed the dynamics of viral fusion. To this end, we forced fusion of TOSV-R18 at the plasma membrane of cells, as described in the bypass assay, but instead measured the increase in fluorescence associated with the dequenching of R18 dye in real-time. In brief, R18-labeled virions were first allowed to attach to A549 cells on ice and viral fusion was then triggered by the addition of buffer at pH ∼5.0. We found that the release of R18 molecules into target cell membranes reached a t_1/2_ of 27 ± 16 sec and was completed 25-50 sec later (Fig. 5G). Almost two-thirds of the plasma membrane-bound virions entered the cells and fused (Fig. 5H). Together, our data indicate that acid-activated penetration of TOSV involves most cell surface-bound virions and is achieved through a fast and efficient fusion process from late endosomal compartments.

### The TOSV envelope glycoprotein Gc is a class-II fusion protein

A growing body of evidence suggests that the Gc envelope glycoprotein of phenuiviruses belongs to the group of class-II membrane fusion proteins (*7*). As a result, phenuiviruses are thought to follow an acid-activated penetration process in line with this class of fusion proteins (*28, 29*). No structural data are currently available for TOSV glycoproteins, but others have resolved the crystal structure of RVFV Gc (*10*). Analysis of the M polypeptide sequence with the blastp algorithm showed that TOSV Gc shares about 48% amino acid identity and 67% similarity with RVFV Gc (Fig. 6A). The fusion domain of RVFV reaches 83% amino acid identity with the corresponding sequence in TOSV Gc. This high degree of conservation suggests a fairly recent evolutionary divergence between TOSV and RVFV.

**Fig. 6:**
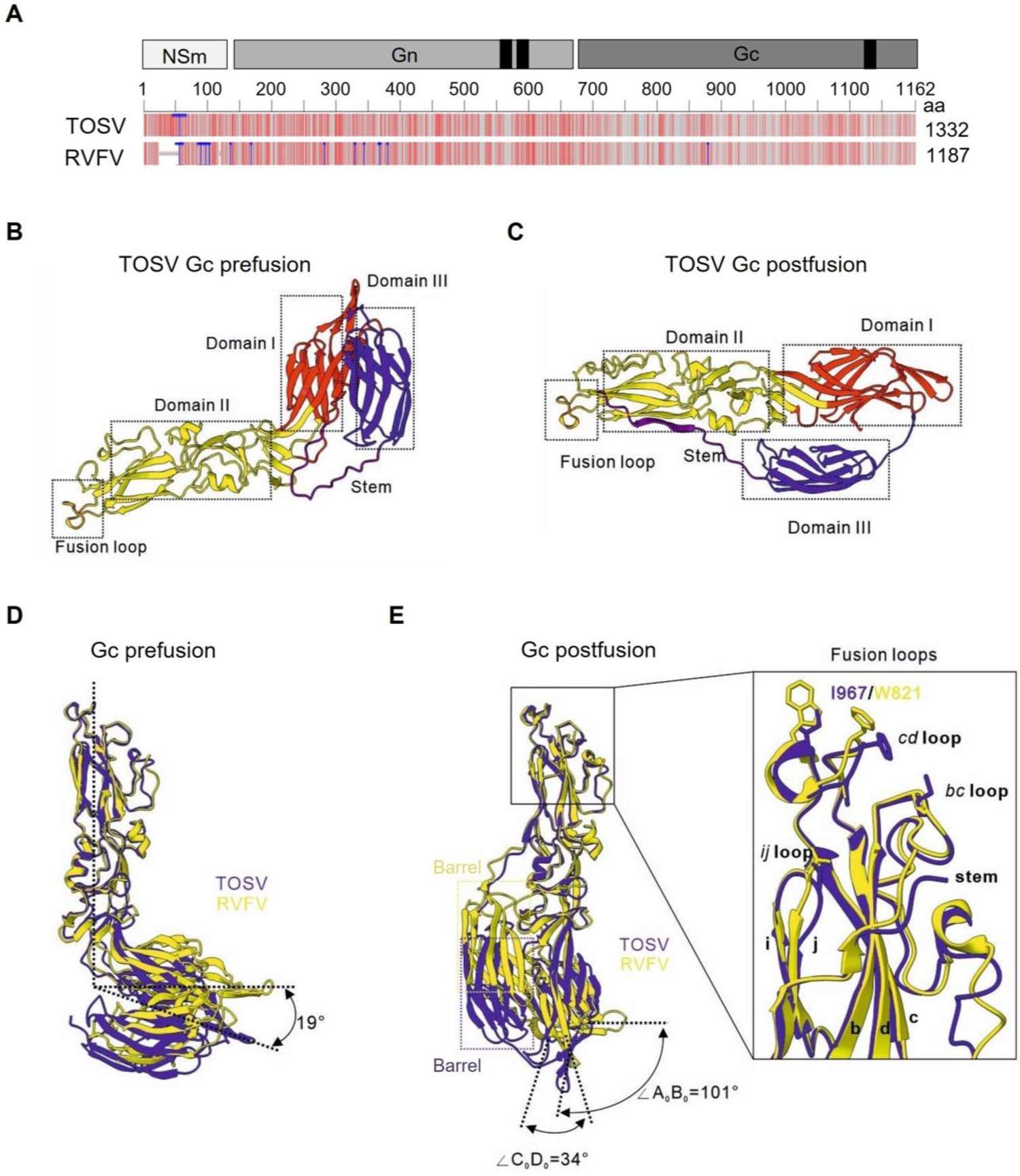
Toscana virus (TOSV) Gc belongs to the group of class-II fusion proteins. (**A**) The M polyproteins of TOSV (strain H4906) and Rift Valley fever virus (RVFV) (strain 35/74) were aligned with blastp suite-2sequences and Multiple Sequence Alignment Viewer. The black boxes indicate the transmembrane domains and the blue lines indicate the insertions into TOSV and RVFV M polyproteins. The grey lines correspond to identical amino acids, the light red lines to similar amino acids and the dark red lines to different amino acids. (**B** and **C**) Pre-(B) and post-fusion (C) conformations of TOSV Gc were predicted using ColabFold, an algorithm primarily based on AlphaFold2, and the Gc structures available for RVFV as modeling models (PDB, 6F9F and 6EGT). The structural predictions were then visualized using PyMOL. Domains I, II and III, typical of class-II membrane fusion proteins with the fusion loop at the end of domain II, appear in red, yellow and blue, respectively. Flexible stem-loops are shown in purple. (**D** and **E**) Pre-(D) and post-fusion (E) conformations of TOSV Gc (blue) were compared with those of RVFV Gc (yellow, PDB, 6F9F and 6EGT) using UCSF ChimeraX. The right box in E shows a magnification of the fusion unit with corresponding loops and amino acids.

The level of identity and similarity in amino acids supports the idea that the Gc glycoproteins of TOSV and RVFV resemble each other structurally. To further test this possibility, we utilized AlphaFold to predict the structure of TOSV Gc ectodomain based on the information available for the RVFV Gc structure. A bent conformation was obtained by comparison with intact RVFV virions (Protein Data Bank [PDB, 6F9F) (*16*), likely corresponding to the canonical pre-fusion orientation of the glycoprotein (Fig. 6B). The Gc AlphaFold prediction exhibited domains I, II and III typical of its phenuivirus orthologs and other viral class-II fusion proteins. The acid-activated conformation of TOSV Gc was predicted using the X-ray post-fusion structure of RVFV Gc (PDB, 6EGT) (*30*) (Fig. 6C). The extent of confidence in the modeling was assessed using an average predicted local distance difference test (pLDDT). The closer the value is to 100, the more likely the prediction is to be close to the real structure. A value below 50 is categorized as low confidence. Both structural models of TOSV Gc achieved a high overall confidence score, with pLDDT values higher than 90 for the individual domains I, II and III.

The AlphaFold pre-fusion conformation of TOSV Gc was virtually identical to the experimentally-resolved pre-fusion structure for RVFV Gc, except for a 19-degree greater angle between domains I and II (Fig. 6D). This difference could be due to the preference of AlphaFold for more energetically stable conformations and not to the original pre-fusion Gc structure itself. The post-fusion Gc forms of the two viruses differed further. The position of A_0_B_0_ and C_0_D_0_ strands in domain I did not undergo major changes after TOSV Gc activation (Fig. 6E and Movie S3). Both A_0_B_0_ and C_0_D_0_ strands had deviations of 101 and 34 degrees, respectively, from the same strands in the RVFV Gc post-fusion structure. A second distinction was the barrel sheet in domain III, which was lower in position for TOSV than for RVFV, and third difference was the presence of an isoleucine residue (I967) in the fusion loop instead of a tryptophan (W821). This latter difference is consistent with the observation made by others that isoleucine at this position allows the distinction between phenuiviruses transmitted by sand flies and ticks and other phenuiviruses (*30*). Overall, our computational approaches showed that TOSV Gc can be confidently considered a viral class-II fusion protein.

### TOSV remains infectious even after acid exposure below the fusion threshold

The activation and priming of class-II viral fusion proteins are described as irreversible steps, and viral fusion proteins act only once (*29*). To check whether TOSV fusion is an irreversible process, we first evaluated the possibility of inactivating the virus, by applying acidic buffers in the absence of target-cell membranes, before infection under neutral-pH conditions. In such an assay, the virus undergoes a transition toward the post-fusion state at the optimal pH. If the transition is irreversible, the fusion protein is no longer able to fuse with target-cell membranes and, thus, the viral particles are rendered noninfectious. Using this approach, we did not observe any negative effect on TOSV infectivity in A549 cells of exposing virions to buffers ranging from pH ∼5.0 to 7.5 for 5 min (Fig. 7A). Infection was even greater when the viral particles were subjected to buffers at pH ∼6.0 and 6.5.

**Fig. 7:**
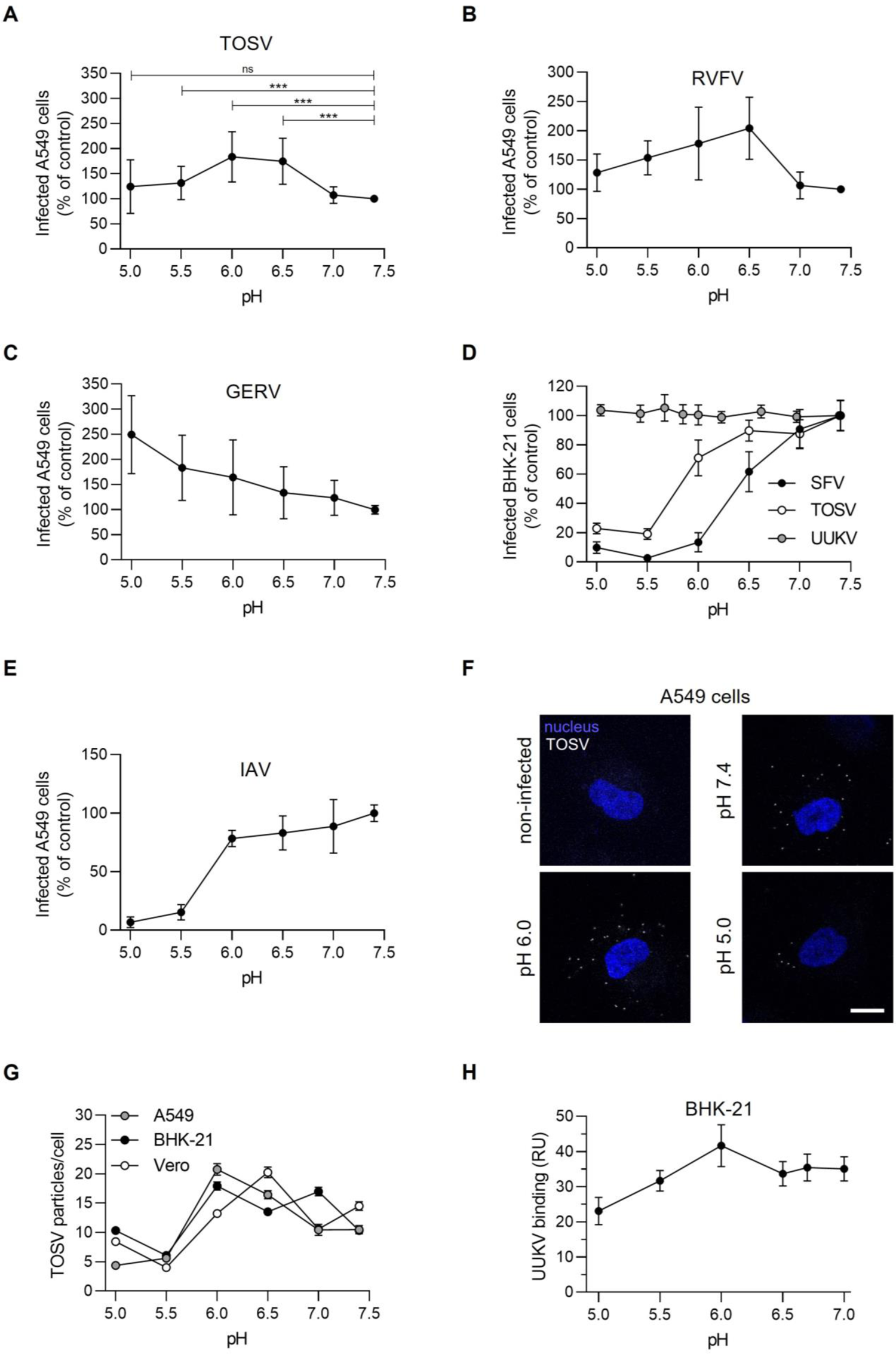
Low pH does not inactivate Toscana virus (TOSV) or other bunyaviruses. (**A**) TOSV was first pretreated at the indicated pH for 5 min at 37 °C. The virus was then neutralized, buffered at pH 7.4 and allowed to infect A549 cells at MOI 2 for 6 h. Infection was quantified by flow cytometry following immunostaining against TOSV structural proteins. Data were normalized to that of samples pretreated at pH 7.4. T-test with Welch’s correction was applied. ***, p<0.001; ns, nonsignificant. (**B** to **E**) A549 cells (B, C and E) and BHK-21 cells (D) were exposed to Rift Valley fever virus (RVFV) construct RVFVΔNSs:EGFP (B), Germiston virus (GERV, C), Semliki Forest virus (SFV) (D), Uukuniemi virus (UUKV, D), TOSV (D) and influenza A virus (IAV, E) pretreated at the indicated pH and analyzed for infection as described in A. (**F**) ATTO647N-TOSV (white) at MOI 1 was pretreated as described in A and then allowed to bind to A549 cells on ice for 90 min before fixation and imaging by confocal microscopy. Nuclei were stained by Hoechst (blue). Scale bar, 10 µm. (**G**) Depicted is the quantification of viral particles bound to A549, BHK-21 and Vero cells as described in F. n > 124 cells. (**H**) Alexa Fluor 647-labelled UUKV was first pretreated at the indicated pH for 5 min at 37 °C, then neutralized with buffer at pH 7.4 and finally allowed to bind to BHK-21 cells at MOI 0.3 on ice for 2 h. Virus binding was quantified by flow cytometry, and data were normalized to samples pretreated at pH 7.0.

To determine whether low pH resistance is a hallmark of TOSV or has larger implications, we expanded our study to related and unrelated viruses with class-I or class-II fusion proteins. Briefly, RVFV, Germiston virus (GERV) and UUKV are bunyaviruses, thus all with a class-II fusion protein and related to TOSV. In addition, SFV is an alphavirus with a class-II membrane fusion protein, and the unrelated influenza A virus (IAV) has a class-I fusion protein (*29*). SFV enters host cells from EEs with a fusion threshold at pH ∼6.2 while the others are late-penetrating viruses (L-PV) with a pH-activation threshold ranging from 5.0 to 6.0 depending on the virus species (*19, 31–34*). Like TOSV, we found that the three other bunyaviruses remained infectious in A549 and BHK-21 cells after exposure to buffers of different pH values (Fig. 7B to 7D). In contrast to bunyaviruses, the infectivity of SFV and IAV was dampened by 90-95% after exposure to pH below 6.0 (Fig. 7D and 7E). We noted that although TOSV infectivity appeared to be lower in BHK-21 cells after exposure to pH ∼5.5 and below, 20-30% of virions remained infectious (Fig. 7D).

To investigate whether the fusion process as such was affected by greater virus binding to cells following exposure to low pH, ATTO647N-TOSV was first subjected to buffers ranging in pH from 5.0 to 7.4. The fluorescently-labeled virions were then allowed to attach to A549, BHK-21 and Vero cells under neutral pH conditions on ice for 1.5 h before imaging (Fig. 7F). Lowering the pH to 6.0 in the binding medium resulted in a higher efficiency of virion attachment to most cell lines, from ∼10 to ∼20 virions per cell (Fig. 7G). Similar results were obtained when UUKV binding to BHK-21 cells was analyzed by flow cytometry (Fig. 7H). More acidic pH probably caused the unmasking of epitopes in viral glycoproteins that promote interactions with one or more cell attachment factors or receptors. The number of cell-bound virions decreased slightly at the lowest pH, *i.e.*, 5-10 virions per cell at pH ∼5.0. Overall, virus binding could not account for enhanced fusion at pH ∼5.5 and below. Combined, the results indicated that TOSV is not inactivated at pH values below the fusion threshold. This suggested that virions remain infectious in late endosomal vacuoles even when the intraluminal acidity was inferior to the fusion threshold.

### Mildly acidic environment primes low-pH-activation of TOSV fusion in LEs

To further examine the effect of pH on TOSV activation and fusion, we forced TOSV-R18 fusion on the A549 cell surface with buffers ranging in pH from 5.0 to 5.8 as described above and monitored the dye dequenching in real-time. Viral fusion could be detected for pH as high as 5.8 and the lower the activation pH, the faster the fusion dynamics (Fig. 8A). In the pH range of 5.0 to 5.5, half of the bound virions had completed fusion within 27-57 sec (Fig. 8B). At pH of 5.8, t_1/2_ was reached within 164 sec, *i.e.*, the process took longer than at lower pH values. When TOSV was first pre-exposed to mildly acidic pH values such as those prevalent in EEs, we found that virions were then able to fuse markedly faster at a pH of 5.8 (Fig. 8C), a pH value typically found in LEs at the beginning of their maturation. The t_1/2_ of the fusion decreased from 168 to 62 sec, which is somewhat similar to the result obtained with an activation pH of 5.5 without pretreatment. Overall, the data demonstrated that TOSV can achieve fusion at higher pHs when exposed to mildly acidic pH for longer periods. The passage through EEs and exposure to a mildly acidic environment most likely favor the activation of TOSV fusion at lower pHs in LEs.

**Fig. 8:**
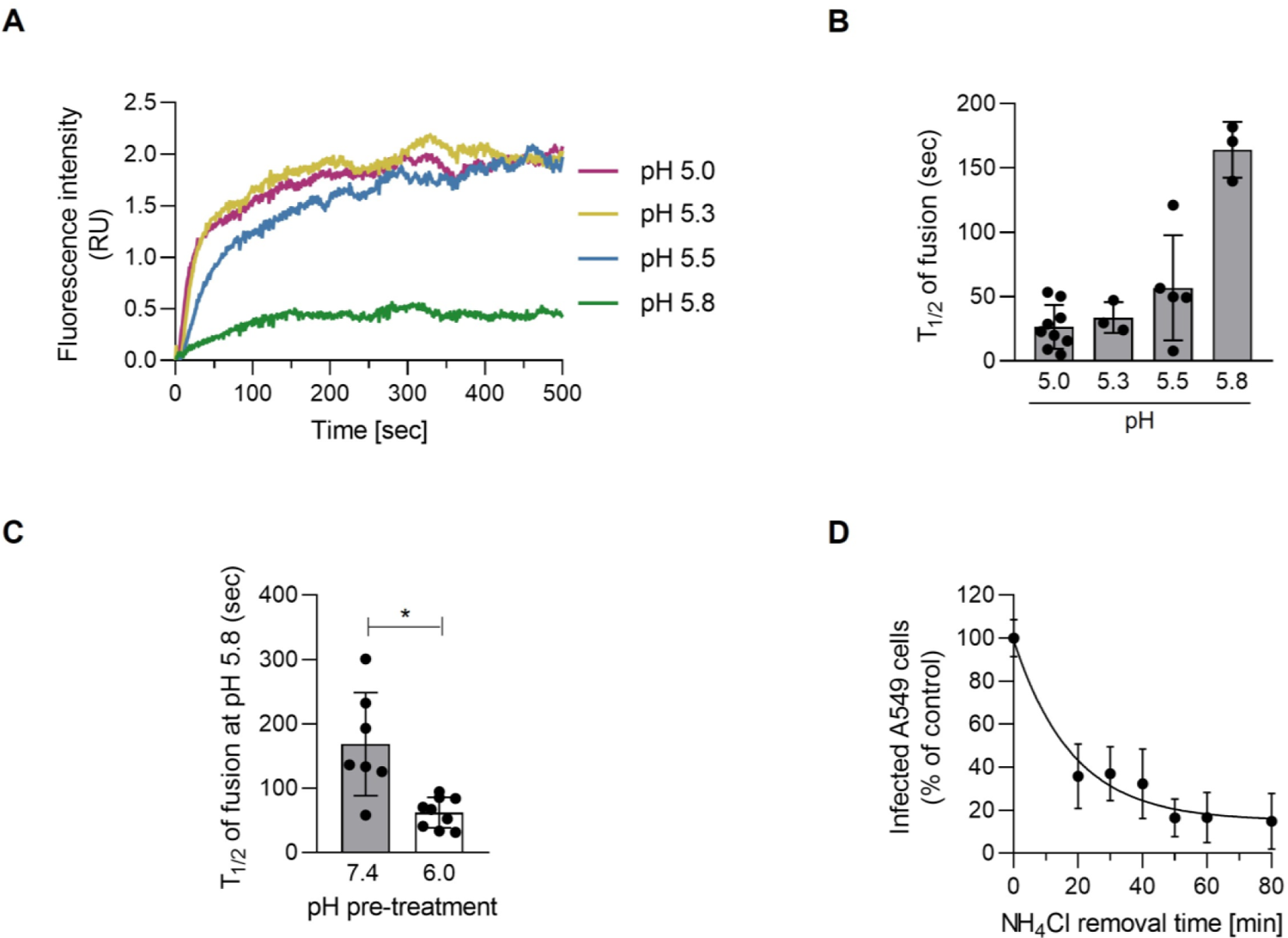
Toscana virus (TOSV) shows remarkable adaptability to the endosomal environment to penetrate cells. (**A**) R18-TOSV binding to A549 cells at MOI 10 was synchronized on ice for 90 min and cells rapidly warmed to 37 °C in a spectrofluorometer. Virus fusion was triggered at the cell surface by adding buffer at the indicated pH, and the fluorescence signal was recorded over 500 sec. The virus fusion-specific R18 release is shown as the result of the values at the indicated pH subtracted from the values corresponding to free diffusion at pH∼7.4. Data were normalized to those at the time point 0. RU, relative unit. (**B**) The half-maximal fluorescence intensity (t_1/2_) was measured in the series of data obtained in A. n > 3. (**C**) R18-TOSV particles were pretreated at pH 6.0 or 7.4 and then neutralized as described in Fig. 7A before being assessed and analyzed as in panel A. The t_1/2_ of fusion was calculated from n > 7. T-test with Welch’s correction was applied. *, p<0.05. (**D**) After the synchronization of TOSV binding at MOI 1 on ice for 90 min, A549 cells were rapidly warmed to 37 °C in the presence of NH_4_Cl (50 mM). NH_4_Cl was then washed out at the indicated times to allow endosomal acidification and the acid-activated penetration of infectious TOSV particles. Samples were harvested 6 h later, and infection was analyzed by flow cytometry. Values were normalized to those from samples for which NH_4_Cl was removed at t0.

### TOSV remains acid-activable in endosomal vesicles for long periods

The resistance of TOSV to low acidity and the ability of virions to fuse at various pHs values led us to postulate that the virus is less prone to inactivation in the endocytic machinery. To examine how long TOSV remains acid-activable in endosomal vesicles, we reversed the approach of adding NH_4_Cl. Virus binding to A549 cells was synchronized on ice, and cells were rapidly warmed in the presence of NH_4_Cl before the weak base was washed out at different times. This assay relies on the fact that the neutralization of endosomal pH by NH_4_Cl is reversible after washing. In other words, we determined the time at which TOSV acid activation was no longer possible. In A549 cells, infection decreased by 60% during the first 30 min and then more slowly until it reached an 80% decrease after 80 min (Fig. 8D). Altogether, these experiments indicated that TOSV infectivity remains high in endosomal vesicles if the virus is not allowed to enter the cytosol after a long period.

Together, our results showed that TOSV resembles late-penetrating viruses in that its entry depends on a normal maturation of LEs. It is transported from EEs to LEs, and its pH of fusion corresponds to that prevailing in LEs. It cannot infect cells in which cargo transport into the degradative branch of the endocytic pathway is blocked. TOSV, and possibly other bunyaviruses, differs remarkedly from other acid-activated viruses in its use of endosomal pH as a cue to enter host cells.

## Discussion

TOSV is a re-emerging human pathogen in southern Europe and northern Africa with more than 250 million people potentially exposed and up to 50% seroprevalence in some areas of the Mediterranean basin (*2*). The TOSV life cycle is however, poorly characterized, and overall, this virus remains neglected. Here, we developed reliable and accurate assays to study the early steps of TOSV infection in both human cell lines and human iPSCs-derived brain cells, which are targeted in the late stages of infection (*35*). We applied flow cytometry, fluorescence microscopy and fluorimetry to analyze each stage of the TOSV entry program, from virus binding and uptake to intracellular trafficking and membrane fusion. To track single viral particles, we labeled TOSV with fluorescent amine-reactive dyes and took advantage of the free amine residues in the viral glycoproteins Gn and Gc. In addition, we relied on the autoquenching property of the lipid dye R18 at high concentrations to examine the acid-activated membrane fusion of virions.

Cryo-electron microscopy images of TOSV showed roughly round particles, homogeneous in size with an average diameter of 103 nm and spike-like projections of 9 nm, quite similar to other phenuiviruses such as UUKV and RVFV (*8, 9*). Our computer-based analysis showed that TOSV Gc shares with RVFV Gc similar molecular weight and 48% homology in amino acid sequence. As expected, the AlphaFold algorithm predicted that Gc structure in TOSV resembles RVFV, with an organization typical of a class-II viral fusion protein. Still, further experimental work is needed to solve the X-ray structure of both TOSV Gn and Gc glycoproteins and to determine whether TOSV particles have the same atypical T=12 symmetry as UUKV and RVFV (*8, 9*).

The binding of TOSV to the cell surface was specific but rather inefficient, *i.e.*, one-quarter of the total input virus. Internalization of virions was completed within 15 min after virus binding and involved about three-quarters of the surface-bound viral particles or less than 20% of the total input virus. In comparison, about 60% of surface-bound viruses fused in intracellular organelles, amounting to nearly 80% of endocytosed virions. One reason for low binding efficiency of the input viral particles to host cells is likely related to the biochemical and biophysical properties of TOSV receptors, the identity of which remains largely undetermined. Similar observations have been made for other phenuiviruses, such as UUKV (*19*). Strikingly, once bound to the cell surface, most viral particles were internalized and trafficked until they reached the appropriate endosomal vesicles to fuse and enter the cytosol.

Confocal microscopy images showed that the number of bound virions per cell was significantly higher than the MOI. This result indicated that a large fraction of viral particles in the virus stocks was noninfectious. Although most surface-bound virions penetrated cells, only one virion every 10 was infectious. Still, the ratio of infectious to noninfectious viral particles was high compared to that of other phenuiviruses, *e.g.*, the ratio in UUKV infectivity is lower than 1:1,000 (*19*). Altogether our results are consistent with a recent study showing that two distinct incomplete phenuivirus populations, which are unable to spread autonomously due to the lack of one or more genome segments, can cooperatively support infection and spread (*36*).

The penetration of enveloped viruses relies on the fusion between the virion envelope and cell membranes. In most cases, fusion is triggered in endosomes after the acid activation of viral glycoproteins. The first observation that TOSV follows the same strategy was the sensitivity of infection to agents that elevate endosomal pH, as is typical for other phenuiviruses (*19, 31, 37*). Using NH_4_Cl, we showed that the first incoming, infectious particles reached the acid-dependent step 5 min after cell warming, and that half had completed this step within 15 min. Another indication was the capacity of cell-bound viruses to fuse to the plasma membrane after exposure to a pH of 5.5 and below. These pH values are consistent with penetration from late endosomal compartments.

From our results, it is furthermore clear that TOSV depends for infection on endocytosis and membrane transport within the classical endocytic machinery. After endocytic uptake, TOSV was observed by confocal microscopy in Rab5a+ EEs. Expression of Rab5a S34N, which inhibits EE maturation and homotypic EE fusion, reduced TOSV intracellular trafficking and infection. Rab5a Q79L, which provokes expansion of EEs and prevents proper LE maturation, also hampered TOSV. Similar observations have been reported for UUKV (*19*). One can conclude that TOSV passes through Rab5a-positive EEs, but to be infectious the virus must reach more acidic downstream organelles, most probably late endosomal compartments.

The hypothesis that the acid-activated step for TOSV occurred in LEs was further supported by the time course and values of pH activation that resemble those of viruses penetrating from LEs such as UUKV and IAV (*38*). These viruses pass the acid-sensitive step typically with a t_1/2_ of 15-20 min and are activated at pH below 6.0. For comparison, viruses fusing in EEs, such as SFV or vesicular stomatitis virus, become NH_4_Cl insensitive within 3-5 min as their internalization is almost instantaneously followed by acid activation (*39*). Other indications were the inhibition of infection observed when LE maturation was hampered at temperatures below 25 °C, after free ubiquitin depletion, and after MT depolymerization. Expression of the inactive mutant Rab7a T22N also impaired TOSV infection. More directly, confocal microscopy coinciding roughly with the time of acid activation showed TOSV in Rab7a+ and LAMP1+ LEs.

Maturing endosomes provide a milieu in which the decreasing pH provides a convenient cue for virus activation (*38*). The TOSV-cell fusion demonstrated that low pH is sufficient to trigger fusion. Proteolytic processing in endosomal vacuoles as observed, *e.g.*, for Ebola virus and SARS-CoV-2 was apparently not needed (*23, 40*). Acid activation occurred in less than 30 sec, consistent with pH-triggered kinetics observed for other acid-dependent viruses (*41*). TOSV most likely follows a fusion process similar to that of RVFV and other phenuiviruses (*16*), the details of which remain to be elucidated to understand what exactly happens in terms of structure of Gn and Gc and mechanisms upon acidification and membrane fusion. More functional investigations will be required to determine whether receptors play a role in these mechanisms.

The fusion process was optimal at a pH below 5.5 but also possible at higher pH despite being delayed by ∼100 seconds. Presumably, TOSV can enter the cytosol from endosomes further upstream in late endocytosis pathways, before reaching appropriate LEs for rapid fusion. In this scenario, TOSV could infect tissues and organs devoid of cellular factors necessary for very late virus penetration. The possibility that the virus penetrates earlier may explain the low efficacy of some perturbants of LE maturation in blocking infection by not only TOSV but also other L-PVs. Typically, drugs such as nocodazole and colcemid interfere with the integrity of MT on which LE depends for maturation (*22*) but have a weak ability to prevent L-PV infection in general (*38*). Nevertheless, whether delayed fusion at suboptimal pH is a specificity of TOSV or a generality among L-PVs remains to be determined.

Another observation supports the notion that infectious entry depends on late endosomal vesicles in their early stages of maturation. The optimal pH for fusion increased from 5.5 to 5.8 when virions were pre-exposed to a mild acidity typical of EEs. Passage through classical EEs was evidently a necessary step for TOSV to reach LEs. Our data indicate that EEs not only play a role in sorting virions in the degradative pathway of the endocytic machinery but are also important in priming the viral fusion that subsequently occurs in later endosomes. These results suggest that Gn and Gc glycoproteins undergo incremental conformational changes during virion trafficking, *i.e.*, before the membrane fusion process itself. Other phenuiviruses and class-II viruses need to be added to this investigation. However, it is tempting to propose a model in which class-II fusion viruses do not depend solely on a narrow pH threshold for acid-activated penetration, but on the progressive decrease of acidity in maturing endosomes. In this model, the pH gradient in the endocytosis pathways would activate multiple successive steps in the viral fusion process as virions travel through the endocytic machinery.

It appears from our data that endosomal acidity triggers viral fusion. However, TOSV, as well as RVFV, UUKV and GERV, remain infectious in the endocytic machinery long after activation. Our results contrast with the widely-accepted view that activation and priming of class-II membrane fusion proteins are irreversible steps and that these fusion proteins act only once (*29*). The fusion of phenuiviruses and some other bunyaviruses may specifically involve intermediate steps that are fully or partially reversible; similar mechanisms have been described for class-III fusion proteins (*42*). The process likely involves other factors than the Gc glycoprotein, such as Gn on the particle surface or proteins and lipids in cellular target membranes. Although probably not a universal approach, as illustrated by the acid-inactivation of SFV, we cannot exclude that some unrelated class-II fusion proteins follow the same strategy. Reports of class-II fusion proteins are essentially based on structural approaches and out-of-cell context assays, whereas TOSV and other bunyaviruses allowed us the real-time analysis of viral fusion only minutes after virion binding and uptake.

Acid-activated membrane fusion and late penetration appear to be features shared by many phenuiviruses and other bunyaviruses (*7, 43, 44*). Our results indicate that TOSV differs from other phenuiviruses in that its penetration seems to rely on LEs in their early stages of maturation, whereas RVFV and UUKV must reach later endosomal compartments and possibly endolysosomes (*19*). TOSV showed great resistance to the degradative branch of the endocytic machinery, remaining infectious for a long time after internalization. The remarkable adaptability of TOSV and other bunyaviruses to endosomal acidity certainly confers to these viruses the advantage of trial and error in endocytosis pathways until they reach endosomes suitable for viral fusion, the detailed structural biology of which remains a challenge for future work. As such, bunyaviruses have likely found a way to expand their possibilities of entering and infecting host cells, and in turn, facilitating their propagation.

## Materials and Methods

### Cell lines

All reagents used for cell culture were obtained from Thermo Fisher Scientific or Merck. Briefly, human A549 and HeLa epithelial cells, HEK293T embryonic kidney cells, Huh-7 hepatocellular carcinoma cells and U87 glioblastoma cells as well as chicken DF-1 embryonic fibroblast cells, murine L929 fibroblast cells, canine MDCK kidney epithelial cells and African green monkey Vero kidney epithelial cells were grown in Dulbecco’s modified Eagle’s medium (DMEM) supplemented with 10% fetal bovine serum (FBS). In addition, 1X non-essential amino acids were added in the culture medium of A549 cells. Baby hamster kidney BHK-21 cells were cultured in Glasgow’s minimal essential medium (GMEM) supplemented with 10% tryptose phosphate broth and 5% FBS. Human Jurkat and SUP-T1 T lymphoblast cells, raji B cells and THP-1 monocyte cells were grown in Roswell Park Memorial Institute (RPMI 1640) containing 10% FBS and the human SH-SY5Y neuroblast cell line in Minimum Essential Medium (MEM)/F12 (Ham’s F12) supplemented with 10% serum. The sand fly cells LLE/LULS40 and LLE/LULS45 were derived from embryos of *Lutzomyia longipalpis* and PPL/LULS49 from *Phlebotomus papatasi*. All sand fly cells were cultured in an L-15-based medium in sealed, flat-sided tubes (Nunc) in ambient air at 28 °C as reported elsewhere (*45, 46*). All cell lines were grown in the presence of 100 units.mL^-1^ penicillin and 100 mg.mL^-1^ streptomycin.

### iPSC-derived neurons

Human induced pluripotent stem cells (iPSCs) derived from a healthy donor (HD6, Heidelberg University) were cultured on Matrigel-coated (Corning) dishes in mTeSR plus medium (STEMCELL Technologies) at 37 °C with 5% CO_2_. Cells were split after 3–5 d, depending on colony size, using EDTA (Sigma). Colonies were scraped off and transferred to Matrigel-coated dishes. The medium was changed every other day. Primary mouse glial cells were prepared as described by Patzke and colleagues (*47*). Briefly, newborn (p0) mouse cortices were isolated and digested with papain for 20 min, cells were dissociated by trituration using a thin pipette tip and passed through a cell strainer. Cells were then plated onto T75 flasks in DMEM supplemented with 10% FBS. Upon reaching confluence, glial cells were trypsinized and re-seeded twice to remove potential trace amounts of mouse neurons before the glial cell cultures were used for co-culture with induced neuron cells. All procedures involving animals were approved by the Governmental Council Karlsruhe, Germany, and were carried out in strict compliance with German Animal Protection Law (TierSCHG) at the Heidelberg University, Germany. Induced human glutamatergic neurons were generated from HD6 iPSCs as previously described (*17*). Briefly, iPSCs were treated with Accutase (Sigma), plated and simultaneously infected with two lentiviruses: one designed to express rtTA driven by the ubiquitin promoter and another one designed to express, in an inducible manner, NGN2 and puromycin driven by the rtTA promoter. One day later, doxycycline was added to the medium at a concentration of 2 µg.mL^-1^ to drive NGN2 and puromycin expression. Two days later, 1 µg.mL^-1^ puromycin was added to the medium during 24h for selection. After selection, the remaining cells were detached with Accutase and re-plated on Matrigel-coated coverslips along with mouse glia. Half of the medium was then changed every second day for eight days, and 2.5% FBS was added to support astrocyte viability. After day 10, induced neurons were cultured in B27/Neurobasal medium containing Glutamax (Gibco) and 5% FBS for a minimum of 21 days before infection with TOSV.

### Viruses

TOSV strain H4906 (lineage B) (*5*), recombinant RVFVΔNSs:EGFP (*48, 49*), GERV (*34*), UUKV strain S23 (*50*), SFV (*32*) and IAV strain PR/8/34 (*51*) have all been described previously.

### Antibodies

Polyclonal antibody against TOSV structural proteins N, Gn and Gc was a generous gift from R.B. Tesh (University of Texas, Galveston, Texas, USA) (*13*). The mouse monoclonal antibody (mAb) against SFV glycoprotein E2 was kindly provided by Prof. Margaret Kielian (Albert Einstein College of Medicine, USA). The polyclonal guinea pig antibody GR1 against N, Gn and Gc structural proteins of GERV was recently described (*34*). The mouse mAb 8B11A3, which targets a linear epitope in the UUKV nucleoprotein N, is a kind gift from Ludwig Institute for Cancer Research (Stockholm, Sweden) (*19*). The mouse mAb that detects the IAV nucleoprotein was purchased from Merck (MAB8257). Secondary antibodies conjugated to AF405 and AF488 were purchased from Molecular Probes.

### Plasmids and reagents

The plasmids encoding Rab5a, Rab7a and LAMP1 wt and mutant molecules tagged with EGFP have all been described previously (*19*). Stock solutions of chloroquine diphosphate (Sigma) and ammonium chloride (NH_4_Cl, Sigma) stocks were prepared in dH_2_O at concentrations of 19 mM and 1 M, respectively. Bafilomycin A1 (BioViotica), colcemid (Cayman Chemical), concanamycin B (BioViotica), MG-132 (Selleck Chemicals), nocodazole (Merck) and taxol (Merck) were all dissolved in 100% dimethyl sulfoxide (DMSO) to prepare stock solutions at 100 µM, 10 mM, 50 µM, 40 mM, 20 mM and 10 mM, respectively. Stock solutions of all drugs were diluted in DMEM at the indicated doses (Figures 1 and 4), which are known not to cause cytotoxicity (*23, 34*). Both dH_2_O and DMSO were included as solvent controls. The hydroxysuccinimidyl (NHS) ester dyes AF488 (Thermo Fisher Scientific) and ATTO647N (Atto-Tec) were dissolved in DMSO (10 and 5 mg.mL^-1^, respectively) while octadecyl rhodamine B chloride (R18, Thermo Fisher Scientific) was dissolved in ethanol (10 mM).

### Virus production, labeling, purification and titration

TOSV, GERV, UUKV and SFV were produced in BHK-21 cells in serum-free medium whereas RVFVΔNSs:EGFP was produced in Vero cells in 2%-containing medium and IAV in MDCK cells in serum-free medium (*13, 23, 34, 51*). All viruses were purified through a sucrose cushion and then titrated. TOSV, GERV, UUKV and SFV were titrated on BHK-21 cells and IAV on MDCK cells using a pfu assay following procedures established in the laboratory (*51, 52*). The titer of RVFVΔNSs:EGFP was determined in A549 cells by quantifying EGFP-positive cells 7 hpi by flow cytometry using a protocol derived from the approach developed by Barrigua and colleagues (*53*). The MOIs are therefore given according to the titers determined in BHK-21 cells for TOSV, GERV, UUKV and SFV, in A549 cells for RVFVΔNSs:EGFP and in MDCK cells for IAV. TOSV was fluorescently labeled using a previously-described method (*52*) in which one and two molecules of AF488 and ATTO647N NHS ester dye, respectively, are conjugated to one molecule of the viral glycoproteins for one of the viral glycoproteins. UUKV was labeled following the same method but with three molecules of AF647 (*14*). Alternatively, TOSV (3×10^9^ pfu.mL^-1^) was labeled with the lipophilic dye R18 (25 µM) (*52*).

### Virus binding and internalization

Virions were allowed to bind to pre-cooled cells in DMEM containing 0.2% BSA and 20 mM 4-(2-hydroxyethyl)-1-piperazineethanesulfonic acid (HEPES) at pH ∼7.4 (binding buffer) on ice for 90 min at indicated MOIs. Where indicated, virions were first buffered to different pH values in 100 mM citric acid (pH < 5.5), 2-(N-morpholino)-ethanesulfonic acid (MES) (5.5 < pH < 6.5), or HEPES (6.5 < pH < 7.4) for 5 min at 37 °C before being returned to pH ∼7.4 and allowed to bind to cells. For internalization assays, cells were rapidly warmed to 37 °C and incubated for the indicated periods. Both virus binding and internalization were analyzed by flow cytometry, confocal microscopy and fluorimetry as described below. To discriminate between internalized and external virions, trypan blue (Sigma) was added to a concentration of 0.01% before the analysis. In flow cytometry- and fluorimetry-based assays, cells were detached from culture plastic by incubation with 0.5 mM EDTA, and virus binding and internalization were performed with cells in suspension in phenol-free binding buffer. In binding competition experiments, cells were first exposed to the indicated amounts of unlabeled TOSV or SFV for 45 min at 4 °C and then to AF488-TOSV at a concentration of 15 nM viral glycoproteins for an additional hour in the presence of unlabeled viruses on ice.

### Infection assay

Cells were exposed to viral particles at the indicated MOIs in the respective medium without serum for 1 h at 37 °C. Virus inoculum was then replaced by the respective complete medium, and infection was allowed for an additional 5 h for TOSV and 7 h for RVFV, GERV, UUKV, SFV and IAV, if not indicated otherwise. Where mentioned, virions were buffered to the indicated pH values as described in the Virus binding and internalization section before the infection of cells. For inhibition experiments, cells were pretreated for 30 min at 37 °C, except for colcemid, nocodazole and taxol. For these three drugs, cells were exposed for 3 h at 4 °C before infection was carried out at 37 °C. For all the drugs, cells were infected in the continuous presence of the inhibitors. To assess the dependence of TOSV entry on temperature, virus entry was first synchronized on ice. Infected cells were then incubated for 50 min at the indicated temperatures before warming to 37 °C in the respective complete medium containing 50 mM NH_4_Cl and buffered with 20 mM HEPES for 6 h. For NH_4_Cl add-in time courses, virus binding to cells was synchronized on ice. Cells were then rapidly warmed to 37 °C, and NH_4_Cl (50 mM) was added at the indicated times. Cells were subsequently incubated at 37 °C and harvested 6-8 h after the initial warm shift. In the reverse approach, virus binding to cells was synchronized on ice, and cells rapidly warmed to 37 °C in the presence of NH_4_Cl (50 mM) before the weak base was washed out at the indicated times. Cells were then incubated for an additional 6 h at 37 °C. Virus infection was monitored by flow cytometry.

### Flow cytometry

Infection was monitored by flow cytometry as previously described (*19*). Briefly, infected cells were fixed and permeabilized by 0.1% saponin before the immunofluorescence staining of newly-produced viral proteins with respective primary antibodies against TOSV, GERV, UUKV, SFV or IAV at concentrations of 2.5 µg.mL^-1^, 1:16,000, 1:1,000, 1:400 and 1:250, respectively. Cells were then washed and exposed to anti-guinea pig or anti-mouse AF405- or AF488-conjugated secondary antibodies (1:500, Thermo Fisher Scientific) for one hour. Alternatively, cells infected with RVFVΔNSs: EGFP were assayed for the EGFP signal, and in binding and internalization experiments, AF488-labeled virions were directly measured. Infection was quantified with a Celesta flow cytometer (Becton Dickinson) and FlowJo software v10.6.2 (TreeStar).

### Viral protein analysis

Purified virus stocks were diluted in lithium dodecyl sulfate (LDS) sample buffer (Thermo Fisher Scientific) and separated by SDS-PAGE (NuPAGE Novex 10% Bis-Tris gel, Thermo Fisher Scientific) as previously described (*5*). Viral proteins were either stained with Coomassie blue or analyzed by fluorography using an LI-COR Odyssey CLx scanner and ImageJ v1.53c [National Institute of Health (NIH), US].

### Cryo-electron microscopy

Sucrose gradient-purified virus particles were washed in a buffer containing 10 mM HEPES, 150 mM NaCl, 1 mM EDTA, pH ∼7.3, pelleted by ultracentrifugation, and fixed with 4% paraformaldehyde. Subsequently, 2.5 µL of the fixed virion solution was applied to degassed Quantifoil R2/2 Cu grids that were discharged at 30 mA for 2 min before sample application. The sample was vitrified in liquid ethane using a Leica EM GP2 plunge freezer at 4 °C and 90% humidity, and sensor blotting from the reverse side for 3 sec. Data were acquired using SerialEM software on a Thermo Fisher Scientific Glacios transmission electron microscope operated at 200 kV and equipped with a Falcon 3 direct electron detector. Before data acquisition, the microscope was adjusted by a comma-free alignment in SerialEM and the gain reference was determined. Regions of interest were identified in low-magnification setups. For high-resolution data acquisition, the nominal magnification was 73,000, resulting in a pixel spacing of 2.019 Å. The camera was operated in linear mode with a dose rate of 16 e-/s/pixel. The total dose was 19.6 e-/Å2 and was divided into 22 dose-fractions that were aligned and gain-corrected in SerialEM. Cryo-EM micrographs were analyzed using ImageJ v1.53c (NIH). The length and width of a viral particle were determined by measuring the largest and smallest distances between peaks in density profile or membranes on the opposite side of the viral particle.

### Fluorescence microscopy

Cells that were exposed to fluorescently labeled virions were mounted with Mowiol (Merck), and if indicated, nuclei were stained with Hoechst 33258 (0.5 μg.mL^-1^, Thermo Fisher Scientific). Live cell imaging was performed in the continuous presence of viral particles. Both fixed and live samples were imaged with a Leica TCS SP8 confocal microscope equipped with an HC PL APO CS2 63x/1.4 N.A. oil immersion objective. In addition, super resolution microscopy was used to image ATTO647N-TOSV mounted in Mowiol on PEI-coated coverslips with a 2-color-STED microscope (Abberior instruments GmbH) as described by Kummer and colleagues (*51*). The STED microscope was equipped with an ×100 Olympus UPlanSApo (NA 1.4) oil immersion objective, and the pixel size was set to 60 nm (confocal) and 15 nm (non-diffracted), respectively. Minor contrast and brightness adjustments of images and Richardson–Lucy deconvolution (regularization parameter of 10^-3^, stopped after 30 iterations) were carried out using Imspector software 16.1.7098 (Abberior instruments). Images were analyzed with ImageJ v1.53c software (NIH) and the Imspector software (Abberior Instruments GmbH).

### DNA transfection

A549 cells (8×10^4^) were transfected with 500 ng of plasmids encoding Rab5a, Rab7a and LAMP1 wt and mutant molecules tagged with EGFP using Lipofectamine 2000 (Thermo Fisher Scientific) according to the manufacturer’s recommendations. Supernatants were replaced by fresh medium 5 h after transfection, and the cells were incubated for an additional 17 h before exposure to TOSV.

### Flow cytometry-based plasma membrane virus fusion

TOSV was forced to fuse with the plasma membrane as previously described (*24*). Briefly, TOSV binding to cells at the indicated MOIs was synchronized on ice, and cells were subsequently exposed to buffers of different pH values as indicated for 90 sec at 37 °C. Infected cells were then washed extensively and incubated in complete medium at pH ∼7.4 supplemented with NH_4_Cl (50 mM) for 7 h. Infection was quantified by flow cytometry following the immunostaining of TOSV structural proteins.

### R18-based virus fusion

The fusion of R18-TOSV with host cell membranes was performed as previously described (*34*). Briefly, cells were detached from the culture surface using 0.5 mM EDTA, and binding of R18-TOSV at MOI 10 to cells in suspension was synchronized on ice in a phenol-free medium at pH ∼7.4 for 90 min. To determine the kinetics of virus penetration, virus-bound cells were rapidly warmed in an FP-8500 fluorometer (Jasco) to 37 °C, and the emission of fluorescence was measured over 90 min. For virus fusion with cell membranes, virus-bound cells were rapidly warmed inside the fluorometer to 37 °C, and the fusion was triggered with buffers of varying pHs as indicated. The fluorescence was measured for 600 sec. Where indicated, virions were pre-exposed to buffers at pH ∼7.4 or 6.0 as described in the Virus binding and internalization section.

### Structural modeling of TOSV Gc

The amino acid sequences of the M segment of TOSV and RVFV, strains H4906 and 35/74, respectively, were first aligned and analyzed with blastp suite-2sequences using a BLOSUM62 matrix and Multiple Sequence Alignment Viewer 1.22.2. ColabFold v1.5.2, an algorithm that combines MMseqs2 with AlphaFold2 (*54*), was then used to predict the structure of TOSV Gc strain H4906 in pre- and post-fusion conformation. For this analysis, the default settings were applied, and the pre- and post-fusion structures of RVFV (PDB, 6F9F (*16*), and PDB, 6EGT (*30*), respectively) served as models. The AlphaFold predictions for TOSV Gc were visualized with the PyMOL Molecular Graphics System, v2.5.4 Schrödinger, LLC. Structural comparisons with RVFV Gc were achieved using UCSF ChimeraX, “matchmaker” plugin to align the models and “morph” plugin to generate conformation change trajectory (*55*). PDB files of TOSV Gc pre- and post-fusion models are available upon request.

### Statistical analysis

Graph plotting and statistics were achieved with Prism v8.0.1 (GraphPad Software). The data shown are representative of at least three independent experiments. Values are presented as the means of triplicate experiments ± standard deviation if not stated differently.

## Acknowledgments

We acknowledge Elodie Chatre and the Imaging Platform Platim, SFR Biosciences, Lyon, as well as Vibor Laketa and the Infectious Diseases Imaging Platform (IDIP) at the Center for Integrative Infectious Disease Research (CIID) Heidelberg. The sand fly cell lines were supplied by the Tick Cell Biobank at the University of Liverpool.

## Author contributions

Conceptualization: JK, PYL Methodology: JK, QX, MO, SK, JC

Investigation: JK, MO, AS, NR, HA, AK, LW, SK

Visualization: JK, QX, PYL

Supervision: JK, ZMU, HGK, FKMS, CA, PYL

Writing – original draft: JK, PYL

Writing – review & editing: JK, QX, MO, AS, HA, SK, ZMU, LBS, FKMS, CA, PYL

## Data and materials availability

All data are available in the main text or the supplementary materials or upon demand within a reasonable time frame.

---

**Fig. S1.**
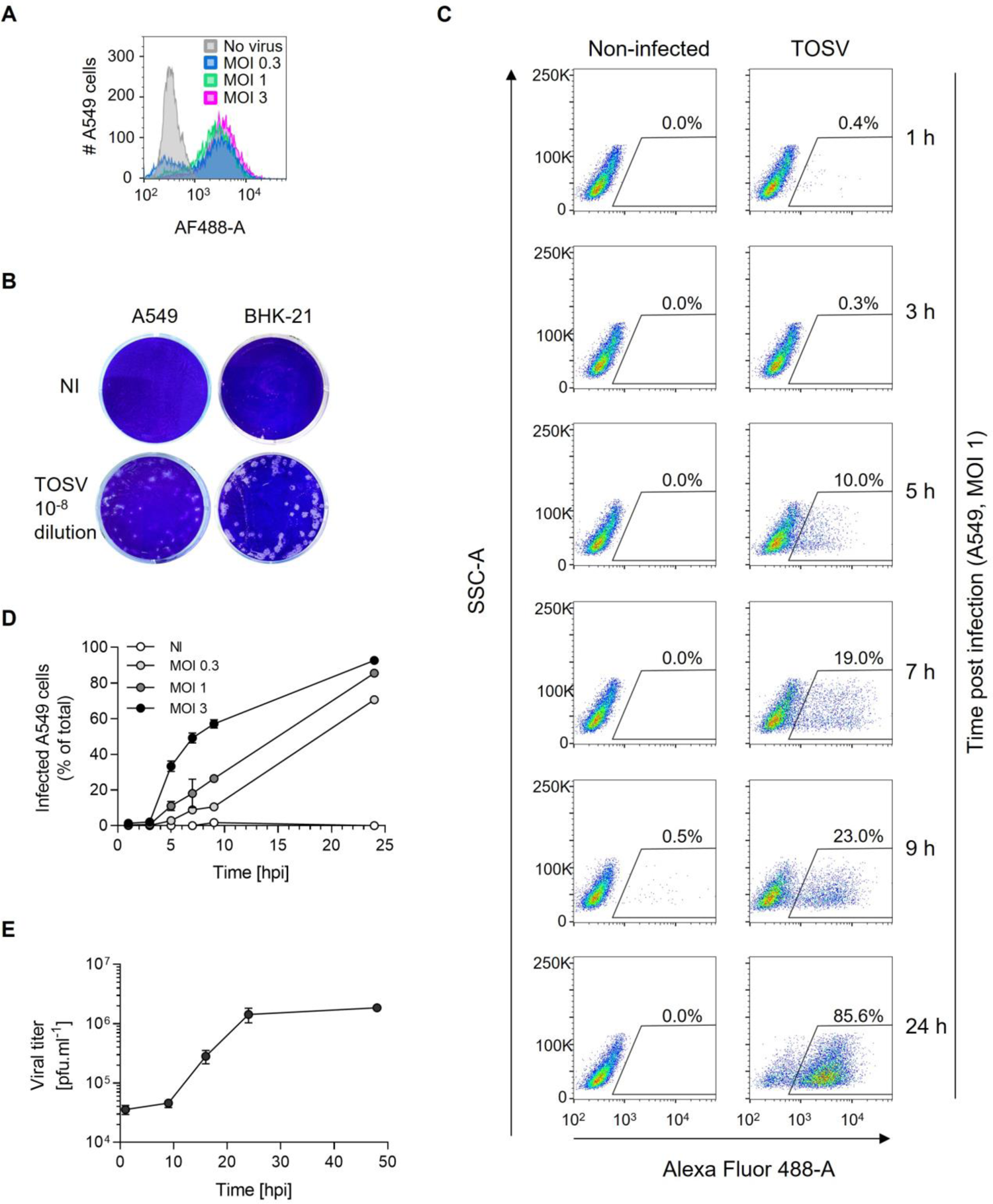
Quantification of Toscana virus (TOSV) infection. (**A**) TOSV was allowed to infect A549 cells at the indicated multiplicities of infection (MOIs) for 24 h. Cells were then fixed and permeabilized, and infection was monitored by flow cytometry after immunostaining against all TOSV structural proteins, i.e., N, Gn and Gc. (**B**) The titer of TOSV stock produced on BHK-21 cells was assessed on A549 cells and BHK-21 cells by a plaque-forming unit (pfu) assay. An example is given here for a 10^-8^ dilution of the virus production. (**C**) A549 cells were exposed to TOSV at MOI 1 for up to 24 h. Infection was monitored by flow cytometry as described in the panel A. (**D**) A549 cells were infected with TOSV at the indicated MOIs over the period of 24 h and analyzed for infection as described in panel C. (**E**) A549 cells were infected with TOSV at MOI 2, and the supernatant from infected cells was collected at time points up to 48 h and analyzed by the pfu assay described in panel B.

**Fig. S2.**
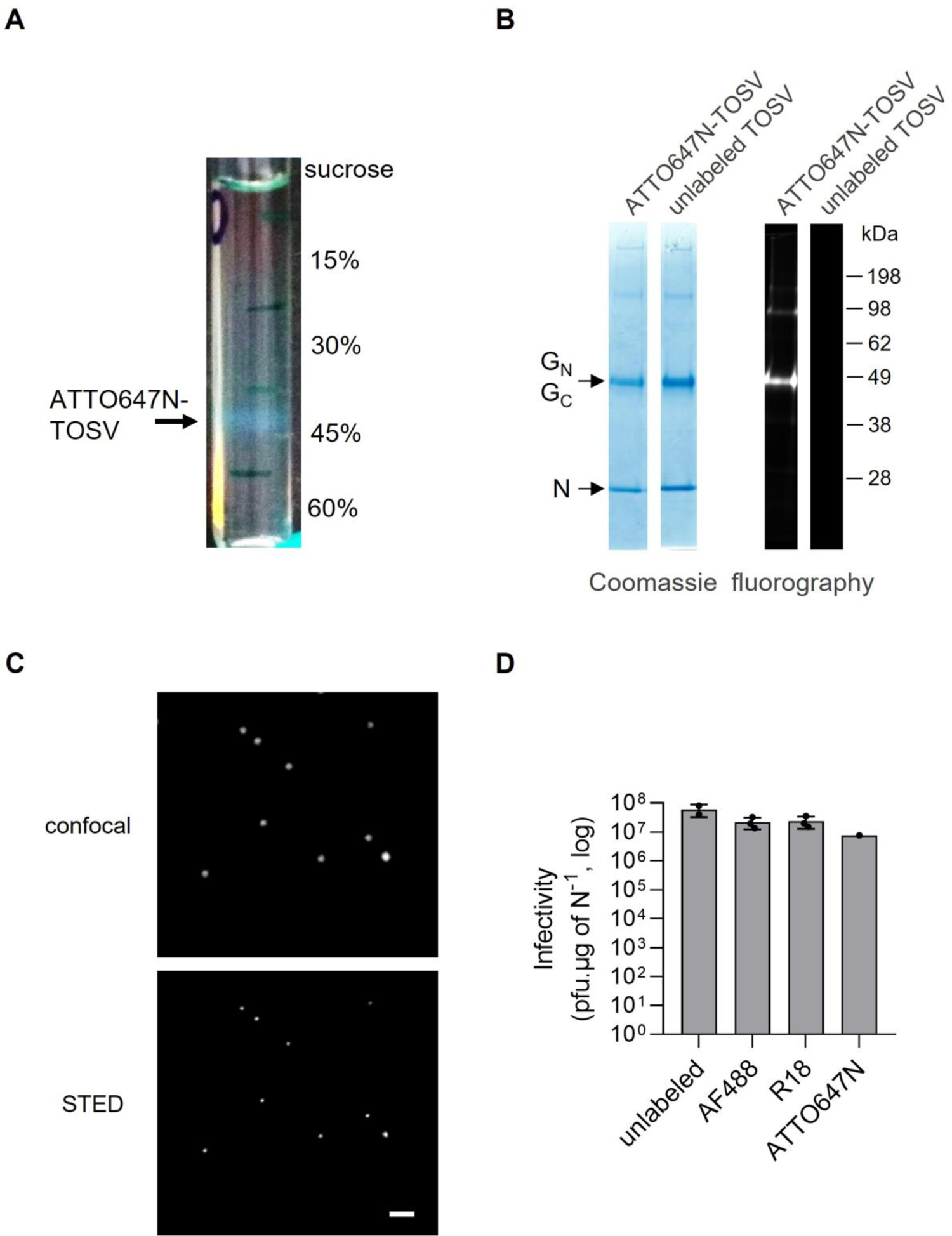
Fluorescence labeling of Toscana virus (TOSV). (**A**) The picture shows a linear sucrose gradient after ultracentrifugation with unbound ATTO647N dye on the top and a band that corresponds to ATTO647N-TOSV particles at a density between 40 and 45% sucrose. (**B**) Fluorescent particles (ATTO647N-TOSV) and unlabeled TOSV were analyzed by nonreducing SDS-PAGE with fluorography (right panel) and then Coomassie blue staining (left panel). (**C**) ATTO647N-TOSV particles were imaged by confocal microscopy (top panel) and STED microscopy (bottom panel). Scale bar, 1 µm. (**D**) Fluorescently labeled TOSV particles were analyzed by the pfu assay shown in Fig. S1B, and the titers were normalized to the amount of the viral nucleoprotein N.

**Table S1.** Susceptibility of different mammalian and sand fly cell lines to Toscana virus (TOSV). **a** Cells were infected with TOSV at MOI 1 for 18 h, fixed, permeabilized and immunofluorescently stained against all TOSV structural proteins. Infection was quantified by flow cytometry, and the sensitivity of cells to TOSV infection (percentage of infected cells) was given as follows: +++ greater than 30%, ++ from 10% to 30%, + from 1% to 10%, - less than 1%. ^b^The production of viral progeny was assessed by pfu titration assay and is given according to the size of plaques 72 hpi as follows: +++ greater than 1 mm, ++ from 0.5 to 1 mm, + less than 0.5 mm, - no plaques. n.d., not determined.

| Cell Line | Species | Tissue | Sensitivity to TOSV infection<sup>a</sup> | Production of new viral particles<sup>b</sup> | Original reference |
| --- | --- | --- | --- | --- | --- |
| A549 | Human | Lung epithelial | +++ | +++ | (1) |
| HEK293T | Human | Embryonic kidney | + | n.d. | (2) |
| HeLa | Human | Cervix epithelial | + | +++ | (3) |
| Huh-7 | Human | Liver epithelial | +++ | ++ | (4) |
| iPSC-derived neurons | Human | Neurons | ++ | n.d. | (5) |
| Jurkat | Human | T lymphoblast | - | n.d. | (6) |
| Raji | Human | B lymphocyte | + | n.d. | (7) |
| SUP-T1<sup>R5</sup> | Human | T lymphoblast | - | n.d. | (8, 9) |
| SH-SY5Y | Human | Neuroblast | +++ | n.d. | (10, 11) |
| THP-1 | Human | Monocyte | - | n.d. | (12) |
| U87<sup>44</sup> | Human | Glial cells | ++ | n.d. | (13-15) |
| BHK-21 | Hamster | Kidney fibroblast | +++ | +++ | (16) |
| DF-1 | Chicken | Embryonic fibroblast | + | ++ | (17) |
| L929 | Mouse | Fibroblast | + | - | (18, 19) |
| MDCK | Dog | Kidney epithelial | + | ++ | (20, 21) |
| Vero E6 | Monkey | Kidney epithelial | +++ | ++ | (22, 23) |
| LLE/LULS40 | Sand fly | Embryonic | + | n.d. | (24) |
| LLE/LULS45 | Sand fly | Embryonic | + | n.d. | (25) |
| PPL/LULS49 | Sand fly | Larva | ++ | n.d. | (25) |

## Legends for Movies S1 to S3

**Movie S1.** Coordinated motion of Toscana virus (TOSV) with Rab5a+ endosomal vesicles. A549 cells stably expressing EGFP-Rab5a were exposed to ATTO647N-TOSV at MOI 10 and imaged every 15 sec at 37 °C by confocal microscopy for 25 min. TOSV particles (magenta) are seen moving with EGFP-Rab5a+ vesicles (green) approximately 11 min post-warming.

**Movie S2.** Coordinated motion of Toscana virus (TOSV) with Rab7a+ endosomal vesicles. A549 cells stably expressing EGFP-Rab7a were exposed to ATTO647N-TOSV at MOI 10 and imaged every 15 sec at 37 °C by confocal microscopy for 25 min. TOSV particles (magenta) are seen moving with EGFP-Rab7a+ vacuoles (green) approximately 24 min post-warming.

**Movie S3.** Conformational transition from the pre-to the post-fusion state of Toscana virus (TOSV) Gc. The structural models of the pre- and post-fusion ectodomains predicted with ColabFold in Figures 6B and 6C were first adjusted for position, and 120 images were then generated to obtain the morph trajectory using UCSF ChimeraX and the “morph” plugin with the wrap parameter set as true. The final movie was recorded at 25 FPS.

